# Time-lagged causal inference corroborates presumed drivers of North Atlantic seabird ecology

**DOI:** 10.64898/2026.07.31.741974

**Authors:** Duncan A. O’Brien, Kate Layton-Matthews, Pol Capdevila, Hannah S. Wauchope, Annette L. Fayet, Tycho Anker-Nilssen, Manuel Ballesteros, Ingar S. Bringsvor, Signe Christensen-Dalsgaard, Nina Dehnhard, Sébastien Descamps, Kjell Einar Erikstad, Kevin Hodges, Svein-Håkon Lorentsen, Tone K. Reiertsen, Vegard Sandøy Bråthen, Hallvard Strøm, Geir Systad, Arnaud Tarroux, Christopher F. Clements

## Abstract

Disentangling causation from correlation is the foundation of the scientific method. Yet, growing evidence suggests that much observational ecology research has not correctly made this distinction due to inappropriate statistical modelling and unappreciated time-delays. Here, we apply time-lagged causal inference techniques to assess the drivers of seabird declines, using multi-decadal North Atlantic seabird data across the behaviour, mass, survival, reproduction and population size of two species’ ecology (Atlantic puffin, *Fratercula arctica*, and black-legged kittiwake, *Rissa tridactyla*). We demonstrate that both climate and anthropogenic activity can suppress breeding success and cause population declines. Moreover, population size is specifically impacted by delayed recruitment effects where both species decline after a lag corresponding to their estimated age of first reproduction. These North Atlantic seabirds are therefore at risk from future environmental and anthropogenic changes, as time-delays may result in populations already on an extinction trajectory prior to changes being detectable in their abundance.

## Introduction

Human alterations to the global biosphere characterise the Anthropocene, threatening the structure and function of Earth systems, and accelerating biodiversity loss and species extinctions (Barnosky *et al*. 2011; Díaz *et al*. 2019). Such losses are believed to be driven by increases in key threats (a.k.a. stressors) which are increasing in line with rates of global consumption (Bellard *et al*. 2012). Typically, the linkage between threats and biodiversity change has emerged from experimental manipulation (Leary & Petchey 2009) supported by observational studies (Capdevila *et al*. 2026), but making statements of causality from ecological observations alone has historically been met with scepticism (Franks *et al*. 2025).

Causal inference techniques (Dee *et al*. 2023; Franks *et al*. 2025) has sought to solve these issues, facilitated by theoretical advancements in econometrics (Pearl 2009; Wooldridge 1999), political science (Kropko & Kubinec 2020) and sociology (Lundberg *et al*. 2021). The key advancement made by such literature is that observational research can reliably capture consequences of stressor changes (Arif & MacNeil 2023; Dee *et al*. 2023) at a fraction of the economic or ethical cost of experiments (and without the simplification of ecological processes), if strong assumptions are fulfilled. However, causal inference design and interpretation requires care as confounding relationships can mask or even reverse true relationships (Arif & MacNeil 2023; Pearl 2009; Vaisey & Miles 2017) if ignored. Many ecological studies are therefore plausibly misinterpreting statistical relationships (Dee *et al*. 2023; Franks *et al*. 2025) due to such confounding, leading to incorrect conclusions and potentially inappropriate solutions to biodiversity loss.

To complicate interpretation further, many observational research fields typically ignore time delays as the naive inclusion of lags can result in biased estimates (Mulder *et al*. 2025; Vaisey & Miles 2017). Unfortunately, in ecological systems, these time delays/lags are a concern given the growing evidence for extinction legacies/’debts’ whereby a species disappears due to events that happened in the past (Hughes *et al*. 2013; Watts *et al*. 2020). Threatened species and ecosystems are particularly vulnerable to such extinction debts (Hughes *et al*. 2013) and highlight how correctly quantifying possible lags is fundamental for their successful protection. Any work assessing threats to biodiversity must therefore carefully identify, measure and fit statistical models considering these potentially confounded and time-delayed relationships to support accurate and timely intervention. Despite this, limited work exists unifying the two fields of causal inference and ecological time lags in real-world ecosystems and populations.

Seabirds represent one of the most threatened vertebrate groups (Croxall et al. 2012) with reported global population declines of up to ∼70% (Paleczny *et al*. 2015). These animals are important to decision makers as they can act as reliable indicators of wider ecosystem change given their upper trophic level position (Cury *et al*. 2011) and existence on the boundary between marine, terrestrial, and atmospheric environments (Mallory *et al*. 2010). Loss of seabirds consequently represents the degradation of one or more ecosystems, so understanding the cause is vital for wider conservation efforts. Extensive research effort has been made to achieve this aim, with long-term monitoring programmes established to understand how different aspects of a seabird population (breeding success, individual birds’ body condition etc.) respond to stress (Mallory *et al*. 2010). Such work has implicated climate change, fishery bycatch, and invasive species in seabird declines (Dias *et al*. 2019), demonstrated that these relatively long lived species can plasticly alter aspects of their ecology to compensate for climatic or fishing stressors (Cairns 1987; Cerini *et al*. 2023), and identified possible time-lagged recruitment effects whereby fledglings only enter the population once they mature and return to the colony to breed (Bird *et al*. 2020; Coulson 2011; Harris & Wanless 2011; Sandvik *et al*. 2012), or resulting from feeding on different aged fish cohorts (Barrett *et al*. 2002; Lewis *et al*. 2001). However, most work is correlative or predictive (Cury *et al*. 2011; Fayet *et al*. 2021; Gibson *et al*. 2023; Layton-Matthews *et al*. 2025; Ponchon *et al*. 2014), rarely reporting the strict assumptions required for causal inference. More specifically, it is the combination of seabirds’ conservation importance, potential confounding arising from plasticity (Grémillet & Charmantier 2010), and lags due to delayed recruitment of both seabirds and their fish-prey, where seabirds feed on certain age-size classes of fish (Barrett 2002; Frederiksen *et al*. 2007), which could create misleading correlative relationships and/or change our understanding of their complex ecology. Seabirds are consequently a prime candidate to explore time-lagged causal inference because of these challenges, alongside the rich theory and data deployed to protect these globally important animals.

This study applies Bayesian structural causal modelling (Pearl 2009) to long-term monitoring data from two North Atlantic seabirds (Atlantic puffin, *Fratercula arctica* & black-legged kittiwake, *Rissa tridactyla* - henceforth puffin and kittiwake) and quantifies the effect of climate and human fishing pressure upon different dimensions of their population ecology. To support these models, we present an analytical workflow that describes our assumptions of the North Atlantic ecosystem, prior to fitting structural causal models that compensate for measurement and time-lag uncertainty. Finally, we propose a robust methodology for estimating the likely time lag of a causal effect.

## Methods

### Defining causal relationships

Before collating data and designing our modelling strategy, we formalised our assumptions of the system and defined our target estimand (i.e. quantitative causal relationship to be investigated - Lundberg *et al*. 2021) under the structural causal model framework (Pearl 2009). Specifically, we have three estimands based upon existing literature for North Atlantic seabirds.

*Estimand 1* - Sea surface temperature (SST) is a causal parent (i.e. has a direct causal effect) of prey fish accessibility (Rose 2005a), and thus becomes an indirect driver of seabird dynamics (Figure 1). Therefore, we hypothesise an increase in SST will increase average population foraging effort *(Fratercula arctica - Fayet et al. 2021)*, and reduce population size *(Uria aalge - Irons et al. 2008; Rissa tridactyla - Sandvik et al. 2014)*, average survival *(Thalasseus maximus - Gibson et al. 2023; F. arctica - Sandvik et al. 2005)*, mass *(Chroicocephalus novaehollandiae scopulinus - Teplitsky et al. 2008)*, and breeding success (multiple species - Layton-Matthews *et al*. 2025; Sydeman *et al*. 2021). We aimed to estimate the average percent change in foraging effort, mass, breeding success, survival and population size in response to a 1°C increase in SST.

**Figure 1.**
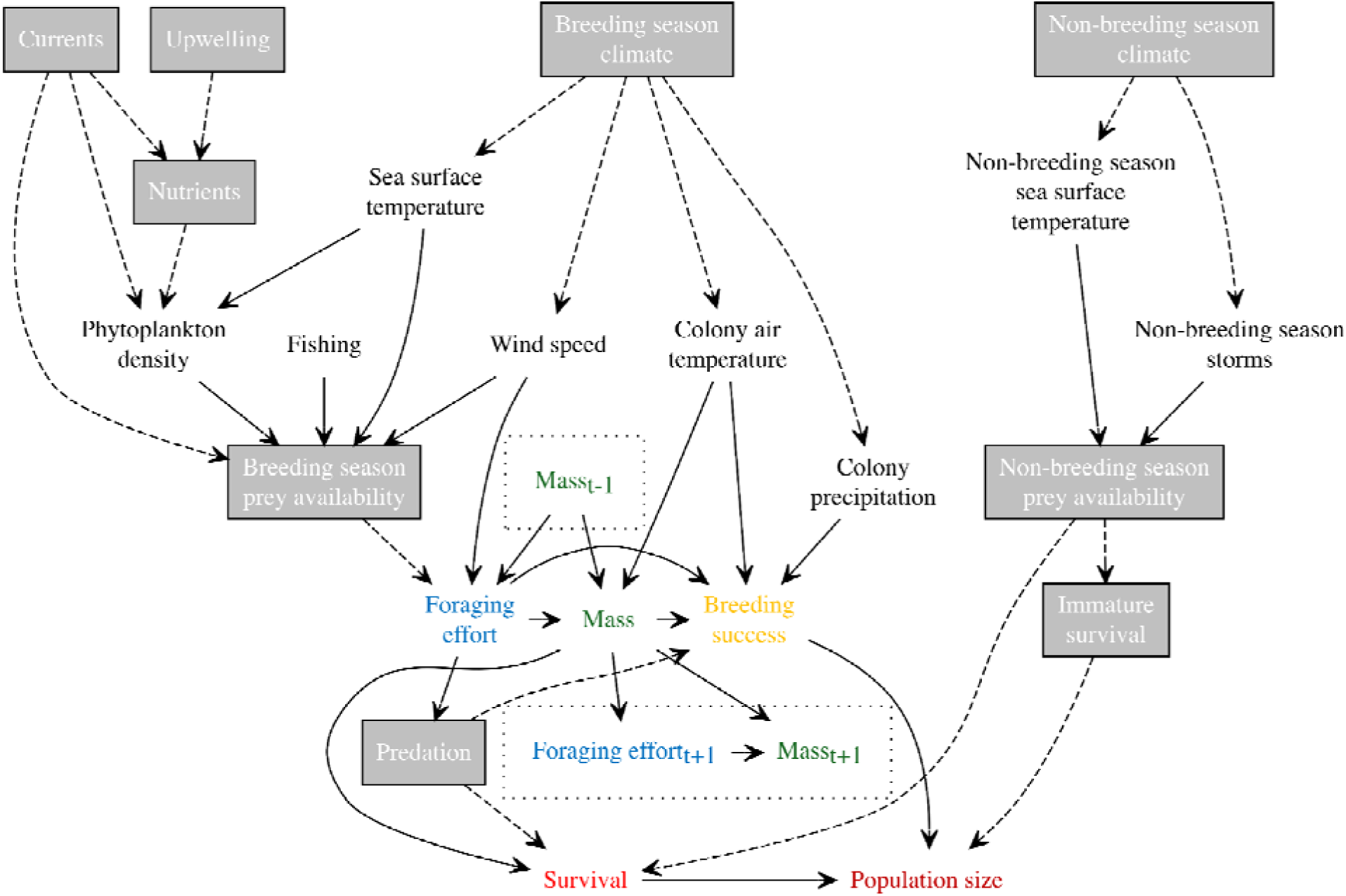
Directed acyclic graph where arrows represent plausible (directional) causal linkages between seabird measurements and extrinsic stressors. Grey nodes represent unmeasured variables. The endogenous influence of time is represented in the *t-1* and *t+1* subscripts.

*Estimand 2* - Fishing pressure is a parent of prey fish accessibility (Gibson *et al*. 2023) and so we hypothesise an increase will cause an increase in average population foraging effort *(Sula variegata - Bertrand et al. 2012)*, and reduce population size *(Phalacrocorax bougainvillii - Barbraud et al. 2018)*, average survivorship *(T. maximus - Gibson et al. 2023)*, mass, and breeding success *(R. tridactyla - Searle et al. 2023)*. We aimed to estimate the average percent change in these variables in response to a unit increase in fishing effort.

*Estimand 3* - Winter storms are a parent to both non-breeding season prey fish accessibility and bird survival (Laurenson *et al*. 2025). We hypothesise an increase in the duration of storms will reduce average survivorship *(F. arctica & U. aalge - Laurenson et al. 2025)* and, as a consequence, population size *(Thalassoica antarctica - Descamps et al. 2015)*. Finally, we aimed to estimate the average percent change in foraging effort, mass, breeding success, survival and population size in response to an additional storm day.

To test our expectations, we applied structural causal model *do-*calculus (Pearl 2009). *do*- calculus links observed data to hypothetical scenarios where a variable has been intervened upon. For example, given a seabird population size *Y* and temperature *X*, we may observe counts across a range of temperatures with probability *P(Y|X)*. However, if we hypothetically intervene and set the entire population’s temperature to *x*, the probability changes to *P(Y|do(X=x)).* Intervening in this way severs all causal parents of *X* as *X* no longer depends on any other variable due to being a constant value (*x*) - and therefore, the association between *do(X=x)* and *Y* is now a causal relationship. While *P(Y|do(X=x))* cannot be estimated directly from observed data, Pearl (2009) demonstrates that we can replicate *do(X=x)* by including an adjustment set *W* that blocks alternative (backdoor) causal paths between *X* and *Y*. If *W* is combined with other assumptions, our estimands can then be calculated from a regression.

To correctly choose *W*, we defined our believed causal relationships in a directed acyclic graph (DAG - Figure 1). It describes a belief that 1) intrinsic population measurements (foraging effort, breeding success etc) influence other intrinsic measurements (Cerini *et al*. 2023), 2) extrinsic climate and anthropogenic stressors act upon different aspects of a population’s biology (Gibson *et al*. 2023; Sandvik *et al*. 2014) and 3) there is a time dependency of intrinsic measurements that flows specifically through seabird morphology/body condition (Cairns 1987 - i.e foraging success will affect future mass and ultimately population size as emaciated individuals die). Moreover, it states that drivers can act through unmeasured variables such as prey availability which open back door paths that may require adjustment sets (*W*) to ensure valid causal inference (Pearl 2009). Justifications for each causal path can be found in Supplementary Material S1.

### Seabird data

Multi-dimensional measurements (behaviour, morphological traits, demographic rates, abundances) have been collected for seabirds at breeding colonies across Norway and Svalbard by the national seabird monitoring programme SEAPOP (www.seapop.no) and tracking program SEATRACK (https://seatrack.net). We focussed on the breeding season of two species - Atlantic puffins and black-legged kittiwakes - breeding at 13 colonies monitored as part of SEAPOP, which together span 1500km of latitude (Supplementary Material S3: Figure S1). Both species are endangered or vulnerable in Europe and have been monitored for more than 20 years, using an annual sampling methodology adapted from international standard methodologies (Supplementary Material S4: Table S1). These sampling strategies target breeding adults as those individuals are easier to observe and/or catch as they are localised to their nests. Descriptions for each of the five measurements used henceforth (**foraging effort, mass**, **breeding success**, **adult survival** and **population size**) can be found in Supplementary Materials S2.

### Exogenous driver data

Information on the drivers of seabird dynamics was collated from three sources - NOAA Daily Optimum Interpolation Sea Surface Temperature (OISST), Norwegian Directorate of Fisheries (Fiskeridirektoratet), and ECMWF Reanalysis v5 (ERA5) - and manipulated in terra v.1.8-54 (Hijmans 2025). Daily atmospheric variable estimates (**wind speed** - m/s 10m above surface, **total precipitation** - mm/day - and **air temperature** - °C 2m above surface) were downloaded from ERA5 (Hersbach *et al*. 2024), whereas daily **sea surface temperature** was downloaded from OISST (Huang *et al*. 2021). Wind speed and sea surface temperature (SST) were extracted for each population’s breeding season kernel and averaged over that population’s breeding season. Air temperature and precipitation were extracted solely at the colonies’ coordinates during the breeding season.

**Fish landings** (tonnes) were downloaded from the Norwegian Directorate of Fisheries, which provides yearly landing information per boat and fishery across major legislative fishing zones in the North Atlantic (www.fiskeridir.no). We subset this dataset to known puffin and kittiwake prey fish - capelin (*Mallotus villosus)*, herring (*Clupea harengus)* and sandeels (*Ammodytes* sp.) (Barrett 2002; Barrett & Krasnov 1996) - and summed the combined landings of these species across fishing zones intersecting with each seabird population’s breeding season kernel. Total landings are known to correlate with fish availability (Maunder & Punt 2004), and we therefore standardised landings by the International Council for the Exploration of the Sea (ICES)’s yearly estimates of spawning stock biomass for the above fish species (ICES 2024). Consequently, our estimate of ‘fishing pressure’ becomes ‘removal of biomass above the system’s capability to recover its fish stocks’, but is not intended to estimate fish availability; only anthropogenic fishing pressure.

Finally, **winter storm** data were estimated from storm tracks generated by the ERA5 project. We identified relevant storms that intersected in a 6 degree radius with the seabird non-breeding season 50% utilisation distributions, with a wind speed >=25m/s (Hodges *et al*. 2011). We then quantified ‘storminess’ as the total duration of storms (days) in November and December. The choice of total storm duration over other characteristics (Reiertsen *et al*. 2021) or principal component analyses (Laurenson *et al*. 2025) gives a more interpretable measure of storm stress, while incorporating both storm frequency and duration in a single variable.

### Obtaining population level estimates from individual level data

In order to compare relationships between behavioural (foraging effort), morphological (body mass), and demographic rates (breeding success and adult survival) of populations, we needed yearly population level estimates for each variable. Breeding success and population size were sampled at the population level, with one value per year per variable. Behavioural, morphological, and survival data were sampled at the individual level (for example, body mass of birds within a population for a given year), and we took annual averages to obtain population level estimates. Due to repeated bird measurements and sexual dimorphism, we did not average the raw data, but rather used hierarchical generalised linear mixture models to produce distributional estimates of population level averages. Full details of the modelling strategy can be found in Supplementary Materials S2.

All models were fit in Stan (Stan Development Team 2025) via its cmdstanr v.0.9.0 (Gabry *et al*. 2025) R v.4.5.1 (R Core Team 2025) interface. Models were run for 2000 iterations over 4 chains, with a warmup of 1000 iterations. This resulted in a total of 4000 draws per model. Diagnostics were assessed for each model to ensure appropriate Rhats, zero divergences, and no maximum treedepth or E-BFMI warnings (Stan Development Team 2025). All distributions reported in this study were constructed from 1000 random draws of the posterior distribution.

### Interpolation of missing years and error propagation

Having obtained annual population level estimates for seabird variables, certain years contained no data due to breaks in monitoring. To estimate missing years, we fit state-space models to each time series and interpolated using the latent states (see Supplementary Materials S2 for exact model specifications). We verified interpolation error by corrupting our complete time series and fitting the same state-space models to our corrupted data as a test dataset. The mean interpolation error was ∼± 0.25 proportions for ordered beta models and ∼± 150 counts for poisson models.

As demonstrated, this data processing workflow (Supplementary Material S3: Figure S2) relies on a sequence of models fit upon previous model outputs. There is consequently a risk that our final estimates may be overly confident/precise if model error is not sufficiently propagated (Simmonds *et al*. 2024). We therefore applied Bayesian posterior fusion (Supplementary Material S2) and drew 1000 draws from this fused posterior to take forward for inference as a time varying representation of population level measurements, and modelling uncertainty. The ultimate time series can be found in Supplementary Material S4: Figure S4-S6.

### Fixed effect panel models

To estimate our estimands, we fit one-way fixed effect models (Kropko & Kubinec 2020), to control for unmeasured time invariant confounding, using brms v2.22.0 (Bürkner 2017) and ordbetareg v0.8.0 (Kubinec 2023). Panel models were fit with different adjustment sets for puffins and kittiwakes as described in Supplementary Materials S2.

This model framework makes a suite of assumptions for our estimands to be a valid causal estimate. Namely, we assume:

1. Our DAG (Figure 1) correctly represents interdependencies between variables.
2. There are no additional population level confounders that vary over time, to those highlighted in the DAG.
3. The conditions a population receives is independent of the outcome (assignment mechanism ignorability) i.e. the SST value is not influenced by the population size.
4. There is no measurement error in the covariates.
5. There is a linear relationship between driver and response (functional form).
6. Given the combination of fixed effects and covariates, the covariance between the hypothesised causal variable and remaining error is zero (strict exogeneity).

Time lags violate the strict exogeneity assumption and to compensate, we included our believed endogeneities in our DAG (Figure 1) but opted to not include all lags in the same model. Instead, we segregated our models by lagged driver effects. This allows us to include lagged variables in the adjustment set where necessary. We then identified the likely lag by supplementing our causal modelling with prediction. Specifically, we identified influential lags by their Bayesian model stacking weights (Yao *et al*. 2017). This procedure involves calculating the Leave-One-Out (LOO) cross-validation for each model and then assigning a weight by minimising the LOO mean squared error (Yao *et al*. 2017). This essentially reframes the process as a model averaging problem, using the predictive ability of each model (i.e. each lag) as a proxy of model quality/importance of lagged effect. This is appropriate because prediction is representative of explanation if the model is causally considered (Scholz & and Bürkner 2025), and the methodology accurately and unbiasedly estimates true relationships as evidenced in Supplementary Material S3: Figure S10. We therefore interpret the lagged causal effect of models which are most strongly represented by the Bayesian stacking weights (i.e. those that contribute more than 33% to predictive ability). Maximum plausible lags were selected for each species using the estimated recruitment time reported by Bird et al. (2020 - 6 years for puffins and 4 years for kittiwakes). While longer lags may occur due to combined recruitment time and seabirds feeding upon 3-year-old fish, extending lags beyond 6 years is unfeasible given the length of the foraging effort and mass time series.

The average causal effect for each lag was then estimated as the ‘*potential increase in response given a one unit increase in driver from its mean, when all other covariates are at their mean, averaged across populations*’ using marginaleffects v0.31.0 (Arel-Bundock *et al*. 2024). As our model predictions can be non-linear across the range of driver values observed (Supplementary Material S3: Figure S11-S13), we chose to report only the effect of increasing the driver one unit over its mean as this estimate represents the imminent future threat to these animals. These changes are large and so we defined a “Region Of Practical Equivalence” (ROPE - Kruschke & Liddell 2018), or a range of causal estimates that are considered equal-to-zero. Specifically, our ROPE was any estimate less than +1% for foraging effort, and greater than -1% for body mass, breeding success, survival and population size as we are fitting a one-sided test and is against these thresholds that we compared our posterior distributions.

Finally, because the error propagation process will increase pseudoreplication and artificially shrink the causal effect uncertainty if unadjusted, we reapplied posterior fusion (Villejo *et al*. 2025) to correctly estimate uncertainty in our average causal effect. This involved fitting a panel model for each draw (see *Interpolation of missing years and error propagation*), estimating the causal effect and LOO cross-validation for each, and combining the causal effects and cross-validations into a ‘global’ estimate. All models reported acceptable diagnostics and were insensitive to our prior choices (priorsense v1.1.0 - Kallioinen *et al*. 2023).

## Results

### Time series dynamics

Most populations declined or persisted at **population sizes** below 1000 (sampled) breeding pairs (Supplementary Material S3: Figure S4 & S5). **Breeding success** was variable (median across all colonies and years [standard deviation], puffin = 0.60 [0.26]; kittiwake = 0.77 [0.40]), with multiple failure years for both species, while **survival** rate was more stable (puffin = 0.86 [0.13]; kittiwake = 0.82 [0.07]). **Body mass** was constant over the monitoring period (puffin = 432.85g [33.74]; kittiwake = 396.20g [24.28]) but the large modelling uncertainty potentially masked nuanced dynamics. Finally, **foraging effort** varied in both magnitude and trend across species and populations: puffins typically foraged between 10 and 30% of the available daylight hours, whereas kittiwakes foraged between 30 and 50%. Extrinsic driver dynamics were similar across all populations though the magnitude varied depending on the colony’s latitude; lower latitude populations (e.g. Anda, Røst) experienced higher SST and precipitation. Fishery pressure was more variable, with populations foraging in the Barents Sea (e.g. Hjelmsøya, Hornøya) most exposed.

### Estimand 1: Sea surface temperature effects

Sea surface temperature is an ubiquitous driver of puffin and kittiwake dynamics (Figure 2, Supplementary Material S4: Table S2), with agreement in the direction of effect across lags. However, certain lags are more influential. These lags may not necessarily be the lag with the largest coefficient estimate as misleadingly strong estimates can emerge from cycles in the time series or cumulative effects, with the dominant lag best explaining patterns in the data.

**Figure 2.**
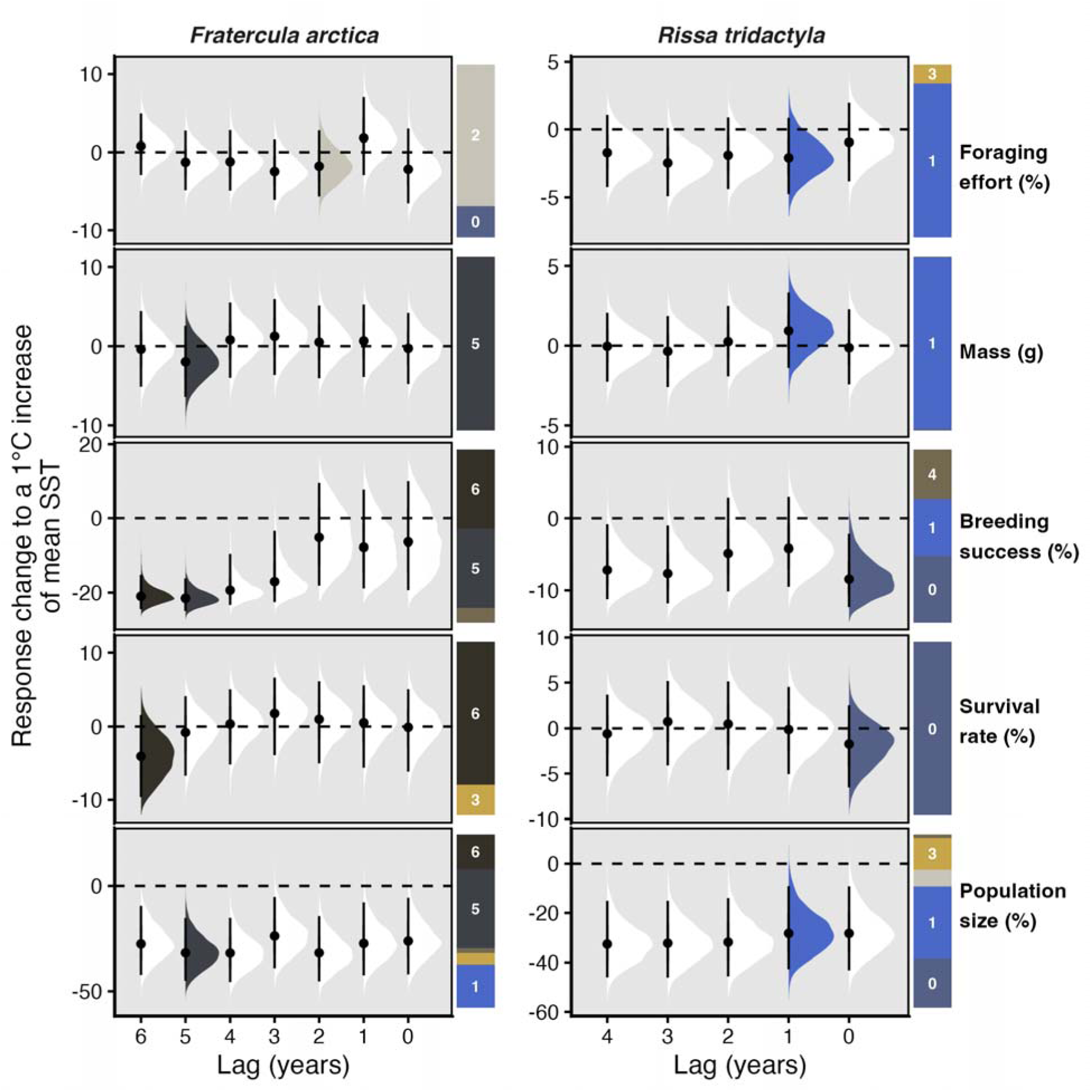
Estimated change in seabird characteristics in response to a 1°C increase in mean sea surface temperature (SST) of their breeding season foraging area across lags. For example, the top two panels represent the % change in foraging effort following a 1°C increase in mean SST, when foraging effort is measured in the same year as SST (X axis = lag0), one year after SST (X axis = lag1) and so on. Estimates are presented as density plots of model posterior distributions where points are median estimates, and error bars the 95% credible interval. Lag estimates are coloured if their contribution to predictive ability exceeds 33%, as estimated by leave-one-out cross validation. The relative importance of each lag is indicated by stacked bar plots to the right of each measurement-SST relationship, whereby each subdivision of the bar is numbered by lag, and its height is that lag’s contribution. The dominant lag may not necessarily be the lag with the largest coefficient estimate. The dashed horizontal line shows the zero-slope.

**Foraging effort** responses do not exceed our ROPE in neither puffins (lag2 median [95% credible interval (Pr(>ROPE))] = -1.76% [-2.89, 2.79 (0.11)]) nor kittiwakes (lag1 = -2.07% [- 4.73, 0.88 (0.02)]). Likewise, **mass** does not change in response to SST (puffin lag5 = -2.01g [- 6.51, 2.65 (0.66)], kittiwake lag2= 0.92g [-1.52, 3.32 (0.29)]). **Breeding success** is negatively impacted by SST with puffins responding approximately at their anticipated recruitment effect lag (lag5 = -21.49% [-24.81, -16.25 (1.00)], lag6 = -20.98% [-24.34, -14.70 (1.00)]), while kittiwakes respond rapidly (lag0 = -8.52% [-12.37, -2.37 (0.99)], lag1 = -4.31% [-9.34, -3.07 (0.83)]). Puffin **survival** responds weakly after a six year lag (lag6 = -3.97% [-9.45, 1.69 (0.83)]) with no detectable effect in kittiwakes (lag0 = -1.79% [-6.71, -2.85 (0.63)]). Finally, **population size** is strongly affected by SST with large declines in puffins’ abundance following a five year lag (lag5 = -31.70% [-44.88, -15.18 (1.00)]) while kittiwakes display a faster response (lag1 = - 28.37% [-42.97, -9.96 (1.00)]).

### Estimand 2: Fishery pressure effects

Fishery pressure differentially impacts different dimensions of seabird ecology (Figure 3 - note a unit change in fishing pressure is on the log scale, Supplementary Material S4: Table S3). **Foraging effort** only weakly increased with fishing pressure in both species (puffin lag1 = 1.10% [-0.21, 2.59 (0.56)]; kittiwake lag1 = 0.82% [-0.48, 2.14 (0.40)]). Only **population size** (puffin lag6 = -6.81% [-9.38, -4.14 (1.00)]; kittiwake lag4 = -9.19% [-13.24, 4.76 (0.1.00)]) and kittiwake **breeding success** (lag1 = -2.40% [-3.89, -0.90 (0.97)], lag4 = -2.60% [-4.10, -1.06 (0.98)]) are driven consistently by fishing pressure.

**Figure 3.**
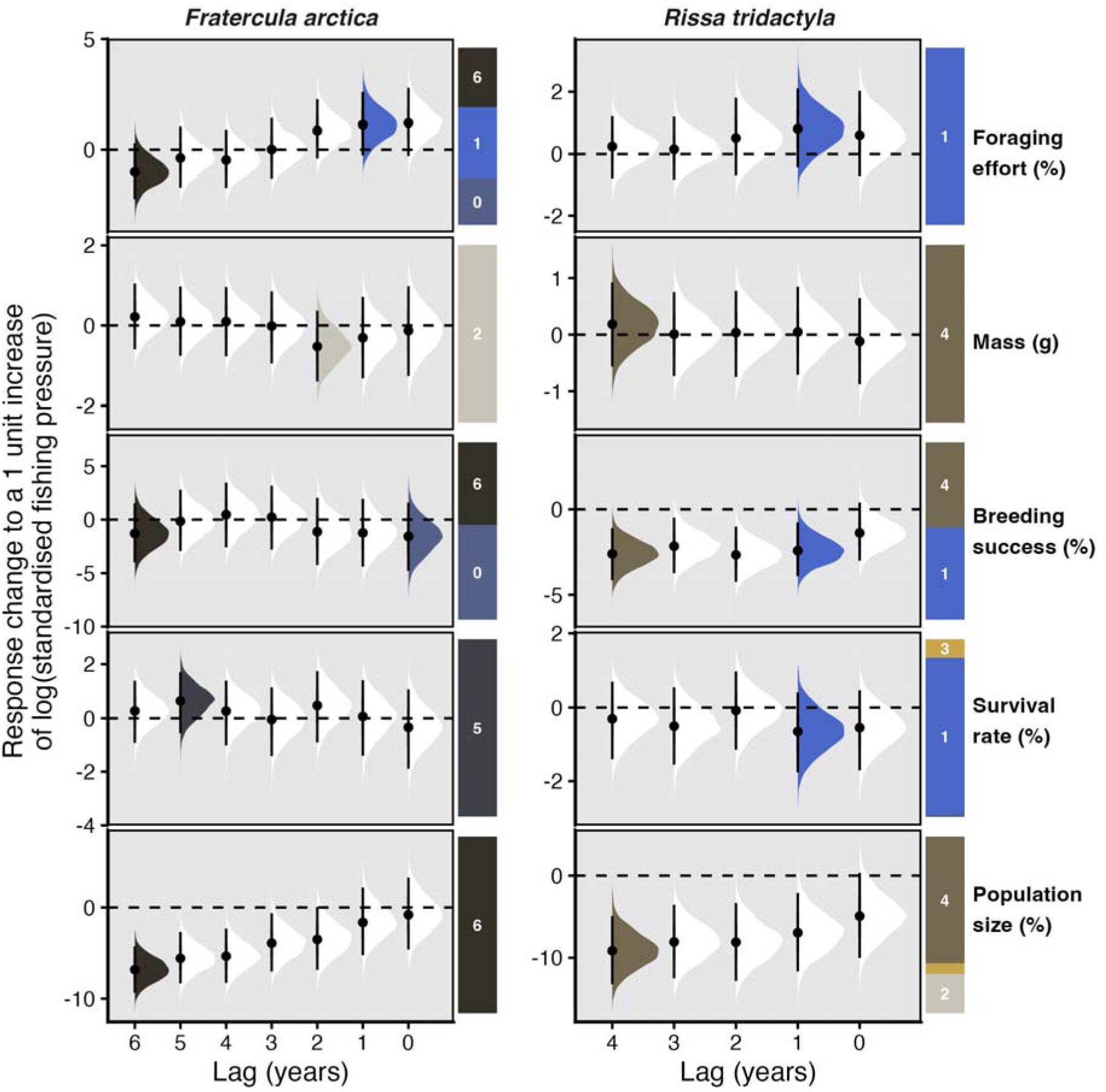
Estimated lagged change in seabird characteristics in response to a one log unit increase in fishing pressure within their breeding season foraging area across lags. Estimates are presented as density plots of model posterior distributions where points are median estimates, and error bars the 95% credible interval. Lag estimates are coloured if their contribution to predictive ability exceeds 33%, as estimated by leave-one-out cross validation. The relative importance of each lag is indicated by stacked bar plots to the right of each measurement-SST relationship, whereby each subdivision of the bar is numbered by lag, and its height is that lag’s contribution. The dominant lag may not necessarily be the lag with the largest coefficient estimate. The dashed horizontal line shows the zero-slope.

### Estimand 3: Storm duration effects

An increase in the number of storm days does not affect seabird ecology (Figure 4; Supplementary Material S4: Table S5) in either species.

**Figure 4.**
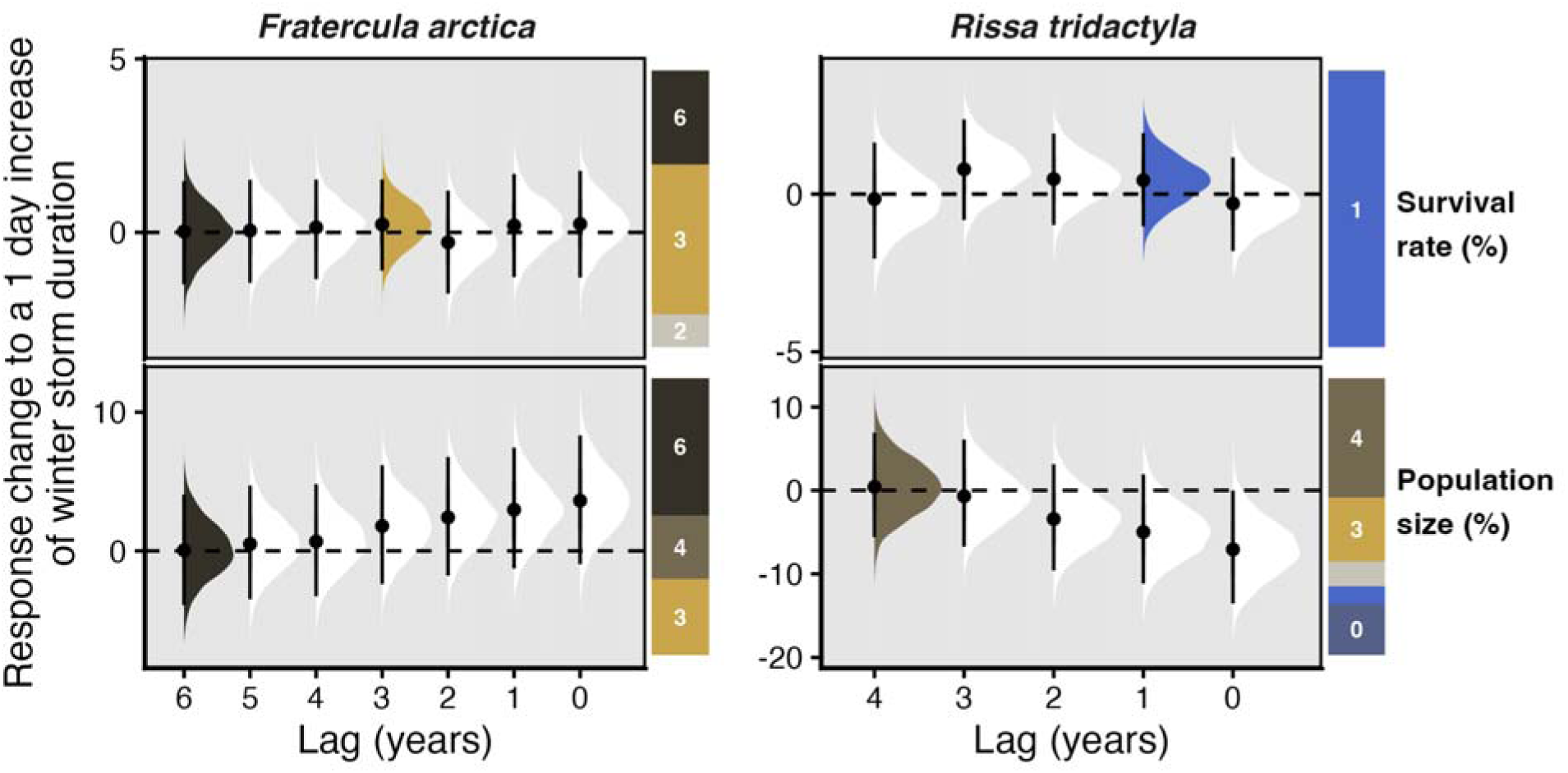
Estimated lagged change in seabird characteristics in response to a 1 day increase in winter storm duration within their non-breeding season foraging area across lags. Estimates are presented as density plots of model posterior distributions where points are median estimates and error bars the 95% credible interval. Lag estimates are coloured if their contribution to predictive ability exceeds 33%, as estimated by leave-one-out cross validation. The relative importance of each lag is indicated by stacked bar plots to the right of each measurement-SST relationship, whereby each subdivision of the bar is numbered by lag, and its height is that lag’s contribution. The dominant lag may not necessarily be the lag with the largest coefficient estimate. The dashed horizontal line shows the zero-slope.

## Discussion

Identifying drivers of biodiversity decline and reliably quantifying their effect(s) is a crucial component of ecological research (Arif & MacNeil 2023; Franks *et al*. 2025) and need of practitioners (Capdevila *et al*. 2026; Hakkinen *et al*. 2022). Causal inference represents the standard for achieving such outcomes from observational data but rarely considers time lags. Using well researched seabird populations, we reiterate their vulnerability to abiotic and anthropogenic stressors (Cury *et al*. 2011; Dias *et al*. 2019) particularly as time delayed effects are present. Such time delayed effects imply both a recruitment effect and potential non-equilibrium behaviour. Early detection of declines is consequently critical as once signals are visible, populations may already be on an extinction trajectory, making reversals slow or even impossible (Hughes *et al*. 2013). The fact that our findings match previous correlative evidence is encouraging given the extensive effort made to mitigate these stressors politically and socially (Hakkinen *et al*. 2022; Klima- og miljødepartementet 2025; OSPAR 2023). However, our identification of lagged effects suggest lags should be explicitly incorporated into decision making (Watts *et al*. 2020).

### Time delayed effects

Using a combination of Bayesian structural causal models (Pearl 2009) and predictive ability (Scholz & and Bürkner 2025), we show that time-lagged effects are estimable in the causal inference framework (Supplementary Material S3: Figure S10). We applied this framework to detect time-lagged effects for North Atlantic puffins and kittiwakes though kittiwakes display shorter lagged responses and respond faster to changes in drivers. Given the SST and fishing pressure drivers explored here primarily act indirectly upon seabirds by depleting prey availability (Cury *et al*. 2011; Reiertsen *et al*. 2014), differences in foraging ecology between the two species is likely the ultimate cause of these species-specific lagged differences. Namely, kittiwakes are surface-foragers (Coulson 2011) and target different fish age classes (Furness & Barrett 1985), potentially restricting their ability to compensate for shifting prey location/depth (Rose 2005b). Kittiwakes may therefore be more vulnerable to prey conditions, which, if insufficient, results in both (breeding) adult and chick mortality. As we do not identify strong effects upon (breeding) adult survival, despite breeding success being reduced, we suggest that the extended lag in puffins corresponds to a recruitment effect (Bird *et al*. 2020) and it is the loss of eggs, chicks and immature birds (Taylor & Gnam 2026) rather than adults that is driving these seabird population declines.

The lagged kittiwake response agrees with previous work (Sandvik *et al*. 2014) which identified population declines following a three-year lag. It is likely that the subtle difference in lag stems from our method of identifying lag importance and challenges in selecting the correct lags to test (Vaisey & Miles 2017). By using predictive ability rather than effect size/significance testing to identify influential lags, we demonstrate a valid method of identifying lagged effects during causal inference, even if multiple lags yield large effect sizes (Supplementary Material S3: Figure S10). For example, lag3 SST has a strong effect on kittiwake population size and some weight (Figure 2), but lag1 is the dominant contributor (i.e. has the most weight).

### Expectation versus observation

Sea surface temperature and fishing pressure remain key drivers of North Atlantic seabird dynamics in our causal analysis. This matches the extensive correlative evidence in the system (Bellard *et al*. 2012; Gibson *et al*. 2023; Irons *et al*. 2008; Sandvik *et al*. 2005, 2012), and strong theory linking both stressors to prey fish availability (Supplementary Material S1). Our findings suggest that population size and breeding success are particularly impacted by prey availability reduction caused by SST increases and fishery pressure. Large-scale climatic changes and fishing actions should therefore be reversed if these populations are to persist at their current colony locations. Fortunately, governmental (Klima- og miljødepartementet 2025) and regional (OSPAR 2023) instruments for the North Atlantic are explicitly targeting these threats via Climate Action Plans and Biodiversity Strategies in a coordinated effort to improve seabird status.

Winter storms are considered an additional threat to seabirds by altering seabirds’ foraging ability during severe weather (Laurenson *et al*. 2025; Reiertsen *et al*. 2021) but we find no evidence for such an effect on the puffin and kittiwake populations assessed. Previous work has used principal component analysis to collapse the complexity of storms (i.e. the wind speed, duration etc) into ‘storminess’ gradients (Laurenson *et al*. 2025) or number of extreme extra- trophic cyclones (Reiertsen *et al*. 2021). Plausibly, our focus on the more interpretable total storm duration is an inappropriate representation of the process acting upon birds, and it is these other storm components that is truly driving the correlative evidence. It is also likely that populations are uniquely vulnerable to storm conditions and by estimating an average treatment effect across a species, we are unable to detect such population-level effects (Laurenson *et al*. 2025; Reiertsen *et al*. 2021). Reiertsen et al. (2021) support this conclusion, only predicting changes in survival in the single population most exposed to extreme storms.

### Assumptions

The validity of our findings are contingent primarily upon how well our directed acyclic graph (DAG) represents reality. There are inevitably linkages that can be debated, such as the order of intrinsic measurements within a breeding season (i.e. behaviour influences morphology, but not vice versa), whether fishing pressure (as defined by quotas and fisher behaviour) is influenced by environmental conditions (Fiorella *et al*. 2021; Golden *et al*. 2024), or that birds respond linearly to these stressors. The DAG presented here (Figure 1) is consequently a transparent demonstration of our beliefs around the North Atlantic system. We welcome discussion around those assumptions and encourage other researchers to use such a tool when designing their own studies.

The use of DAGs also provides a means to formalise, and compensate for, possible confounding caused by missing data (Mohan & Pearl 2014). For example, as breeding adults are almost solely the focus of national seabird monitoring programs, we have necessarily focussed on a sub-sample of the entire seabird population. Our DAG makes this relationship explicit in the non-breeding season via the inclusion of immature survival, while, in the breeding season, it argues that adult birds are missing-at-random (Mohan & Pearl 2014) because the threats themselves remove non-breeding adults from the dataset. This scenario results in the causal effect estimates being relevant to the entire bird population, not just breeding adults as we must assume for other correlative studies.

Despite this, we acknowledge that we have focussed solely on averages despite it being clear that variability in individual bird responses and spatial environmental conditions is also influential on seabird ecology (Mckendrick *et al*. 2026; Wood *et al*. 2021). By averaging across individual birds, core-area kernels, and days within a breeding season, our ‘average treatment effect’ (ATE) estimate potentially smooths out colony or individual level signals and shrinks the estimate towards zero. Repeating these analyses at the individual level (i.e. for foraging effort, mass and survival if tracked across the same birds) would alternatively allow estimation of Average Treatment Effect on the Treated (Lundberg *et al*. 2021) for stressed vs unstressed birds, but such datasets are limited (Mallory *et al*. 2010).

## Conclusion

Overall, our results highlight that: (1) time-lags can reliably be integrated into causal inference analyses, (2) consideration of time-lagged effects sheds light on both the timescales and possible mechanisms of seabird population change; and (3) observational causal inference techniques generally support conventional wisdom around North Atlantic seabird ecology. Due to the challenges involved, this study represents a holistic assessment of North Atlantic seabirds, combining long-term monitoring data across the breadth of seabird ecology, climate models, error propagation, time lags, and observational causal inference to identify drivers of seabird ecology. Fishing pressure on some important prey-fish stocks for seabirds does influence seabird populations but broader climatic changes are more impactful in both magnitude and persistence, leading both species at risk (IUCN 2022; Sandvik *et al*. 2014).

## Data availability statement

Raw tracking data is available following a data request from SEATRACK (https://seatrack.net) whereas SEAPOP data is available from the NINA data portal (https://www2.nina.no/seapop/seapophtml/). ERA5 atmospheric data (air temperature, wind speed, and precipitation) can be found through the Copernicus Climate Change Service Climate Data Store (https://doi.org/10.24381/cds.4991cf48), while OISST data is available from the NOAA National Centers for Environmental Information (https://www.ncei.noaa.gov/products/optimum-interpolation-sst). Fishing data was extracted from Fiskeridirektoratet published reports (www.fiskeridir.no) and is provided alongside storm data in the associated code repository. All code and models are available in the public repository: https://github.com/duncanobrien/seabird-causal.

## Acknowledgments

This study was supported by NERC grant NE/Z000130/. Data collected through the SEAPOP programme (www.seapop.no/en) was financed by the Norwegian Environment Agency, the Norwegian Ministry of Energy via the Research Council of Norway (Grant number 192141) and Offshore Norge. The SEATRACK programme (https://seatrack.net) is funded by the Norwegian Ministry of Climate and Environment, the Norwegian Ministry of Energy via the Research Council of Norway, the Norwegian Environment Agency, the Norwegian Coastal Administration and Offshore Norge along with 16 energy companies. Data were also collected by the Norwegian Polar Institute (via the Monitoring of Svalbard and Jan Mayen (MOSJ) program, www.npolar.mosj.no). Data collected prior to SEATRACK and SEAPOP received financial support from the preceding institutions of the Norwegian Environment Agency and Equinor, as well as the Research Council of Norway (Hornøya, Grant number 216547), the Norwegian Polar Institute and the County Governors of Nordland, Troms, and Finnmark. Permits to work in the colonies were obtained from the local County Governors, with permissions to handle birds and equip them with loggers granted by the Norwegian Environment Agency and the Norwegian Food Safety Authority. We thank all those responsible for the seabird tracking and monitoring at the different sites and their numerous field assistants and students who contributed to the data collection over the years. Finally, we thank the Norwegian Coastal Administration for facilitating stays at Sklinna and Anda lighthouses, and Hornøyas venner for Hornøya lighthouse.

## References

Arel-Bundock, V., Greifer, N. & Heiss, A. (2024). How to interpret statistical models using marginaleffects for R and python. J. Stat. Soft., 111, 1–32.

Arif, S. & MacNeil, M.A. (2023). Applying the structural causal model framework for observational causal inference in ecology. Ecological Monographs, 93, e1554.

Barbraud, C., Bertrand, A., Bouchón, M., Chaigneau, A., Delord, K., Demarcq, H., et al. (2018). Density dependence, prey accessibility and prey depletion by fisheries drive Peruvian seabird population dynamics. Ecography, 41, 1092–1102.

Barnosky, A.D., Matzke, N., Tomiya, S., Wogan, G.O.U., Swartz, B., Quental, T.B., et al. (2011). Has the Earth’s sixth mass extinction already arrived? Nature, 471, 51–57.

Barrett, R.T. (2002). Atlantic puffin Fratercula arctica and common guillemot Uria aalge chick diet and growth as indicators of fish stocks in the Barents Sea. Mar Ecol Prog Ser, 230, 275–287.

Barrett, R.T., Anker-Nilssen, T., Gabrielsen, G.W. & Chapdelaine, G. (2002). Food consumption by seabirds in Norwegian waters. ICES Journal of Marine Science, 59, 43–57.

Barrett, R.T. & Krasnov, Y.V. (1996). Recent responses to changes in stocks of prey species by seabirds breeding in the southern Barents Sea. ICES Journal of Marine Science, 53, 713–722.

Bellard, C., Bertelsmeier, C., Leadley, P., Thuiller, W. & Courchamp, F. (2012). Impacts of climate change on the future of biodiversity. Ecology Letters, 15, 365–377.

Bertrand, S., Joo, R., Arbulu Smet, C., Tremblay, Y., Barbraud, C. & Weimerskirch, H. (2012). Local depletion by a fishery can affect seabird foraging. Journal of Applied Ecology, 49, 1168–1177.

Bird, J.P., Martin, R., Akçakaya, H.R., Gilroy, J., Burfield, I.J., Garnett, S.T., et al. (2020). Generation lengths of the world’s birds and their implications for extinction risk. Conserv Biol, 34, 1252–1261.

Bürkner, P.-C. (2017). brms: An R package for Bayesian multilevel models using Stan. Journal of Statistical Software, 80, 1–28.

Cairns, D.K. (1987). Seabirds as indicators of marine food supplies. Biological Oceanography, 5, 261–271.

Capdevila, P., O’Brien, D., Marconi, V., Johnson, T.F., Freeman, R., McRae, L., et al. (2026). Halting predicted vertebrate declines requires tackling multiple drivers of biodiversity loss. Science Advances, 12, eadx7973.

Cerini, F., Childs, D.Z. & Clements, C.F. (2023). A predictive timeline of wildlife population collapse. Nature Ecology & Evolution, 7, 320–331.

Coulson, J. (2011). The Kittiwake. Poyser Monographs. 1st edn. Bloomsbury Publishing.

Cury, P.M., Boyd, I.L., Bonhommeau, S., Anker-Nilssen, T., Crawford, R.J.M., Furness, R.W., et al. (2011). Global seabird response to forage fish depletion—one-third for the birds. Science, 334, 1703–1706.

Dee, L.E., Ferraro, P.J., Severen, C.N., Kimmel, K.A., Borer, E.T., Byrnes, J.E.K., et al. (2023). Clarifying the effect of biodiversity on productivity in natural ecosystems with longitudinal data and methods for causal inference. Nature Communications, 14, 2607.

Descamps, S., Tarroux, A., Varpe, Ø., Yoccoz, N.G., Tveraa, T. & Lorentsen, S.-H. (2015). Demographic effects of extreme weather events: snow storms, breeding success, and population growth rate in a long-lived Antarctic seabird. Ecology and Evolution, 5, 314– 325.

Dias, M.P., Martin, R., Pearmain, E.J., Burfield, I.J., Small, C., Phillips, R.A., et al. (2019). Threats to seabirds: A global assessment. Biological Conservation, 237, 525–537.

Díaz, S., Settele, J., Brondízio, E.S., Ngo, H.T., Agard, J., Arneth, A., et al. (2019). Pervasive human-driven decline of life on Earth points to the need for transformative change. Science, 366, eaax3100.

Fayet, A.L., Clucas, G.V., Anker-Nilssen, T., Syposz, M. & Hansen, E.S. (2021). Local prey shortages drive foraging costs and breeding success in a declining seabird, the Atlantic puffin. Journal of Animal Ecology, 90, 1152–1164.

Fiorella, K.J., Bageant, E.R., Schwartz, N.B., Thilsted, S.H. & Barrett, C.B. (2021). Fishers’ response to temperature change reveals the importance of integrating human behavior in climate change analysis. Science Advances, 7, eabc7425.

Franks, D.W., Ruxton, G.D. & Sherratt, T. (2025). Ecology needs a causal overhaul. Biological Reviews, 100, 1950–1969.

Frederiksen, M., Edwards, M., Mavor, R. & Wanless, S. (2007). Regional and annual variation in black-legged kittiwake breeding productivity is related to sea surface temperature. Mar Ecol Prog Ser, 350, 137–143.

Furness, R.W. & Barrett, R.T. (1985). The food requirements and ecological relationships of a seabird community in North Norway. Ornis Scandinavica (Scandinavian Journal of Ornithology), 16, 305–313.

Gabry, J., Češnovar, R., Johnson, A. & Bronder, S. (2025). cmdstanr: R Interface to “CmdStan” (manual).

Gibson, D., Riecke, T.V., Catlin, D.H., Hunt, K.L., Weithman, C.E., Koons, D.N., et al. (2023). Climate change and commercial fishing practices codetermine survival of a long-lived seabird. Global Change Biology, 29, 324–340.

Golden, A.S., Baskett, M.L., Holland, D., Levine, A., Mills, K. & Essington, T. (2024). Climate adaptation depends on rebalancing flexibility and rigidity in US fisheries management. ICES Journal of Marine Science, 81, 252–259.

Grémillet, D. & Charmantier, A. (2010). Shifts in phenotypic plasticity constrain the value of seabirds as ecological indicators of marine ecosystems. Ecological Applications, 20, 1498–1503.

Hakkinen, H., Petrovan, S.O., Sutherland, W.J., Dias, M.P., Ameca, E.I., Oppel, S., et al. (2022). Linking climate change vulnerability research and evidence on conservation action effectiveness to safeguard European seabird populations. Journal of Applied Ecology, 59, 1178–1186.

Harris, M.P. & Wanless, S. (2011). The puffin. A&C Black.

Hersbach, H., Comyn-Platt, E., Bell, B., Berrisford, P., Biavati, H., Horányi, A., et al. (2024). ERA5 post-processed daily statistics on single levels from 1940 to present.

Hijmans, R.J. (2025). terra: Spatial data analysis (manual).

Hodges, K.I., Lee, R.W. & Bengtsson, L. (2011). A comparison of extratropical cyclones in recent reanalyses ERA-interim, NASA MERRA, NCEP CFSR, and JRA-25. Journal of Climate, 24, 4888–4906.

Huang, B., Liu, C., Banzon, V., Freeman, E., Graham, G., Hankins, B., et al. (2021). Improvements of the Daily Optimum Interpolation Sea Surface Temperature (DOISST) Version 2.1. Journal of Climate, 34, 2923–2939.

Hughes, T.P., Linares, C., Dakos, V., van de Leemput, I.A. & van Nes, E.H. (2013). Living dangerously on borrowed time during slow, unrecognized regime shifts. Trends in Ecology & Evolution, 28, 149–155.

ICES. (2024). Working group on the integrated assessments of the norwegian sea (WGINOR; outputs from 2023 meeting).

Irons, D.B., Anker-Nilssen, T., Gaston, A., Byrd, G.V., Falk, K., Gilchrist, G., et al. (2008). Fluctuations in circumpolar seabird populations linked to climate oscillations. Global Change Biology, 14, 1455–1463.

IUCN. (2022). The IUCN Red List of Threatened Species.

Kallioinen, N., Paananen, T., Bürkner, P.-C. & Vehtari, A. (2023). Detecting and diagnosing prior and likelihood sensitivity with power-scaling. Statistics and Computing, 34, 57.

Klimaog miljødepartementet. (2025). Nasjonal handlingsplan for å bedre situasjonen for sjøfuglbestandene 2025–2035.

Kropko, J. & Kubinec, R. (2020). Interpretation and identification of within-unit and cross-sectional variation in panel data models. PLOS ONE, 15, e0231349.

Kruschke, J.K. & Liddell, T.M. (2018). The Bayesian New Statistics: Hypothesis testing, estimation, meta-analysis, and power analysis from a Bayesian perspective. Psychonomic Bulletin & Review, 25, 178–206.

Kubinec, R. (2023). Ordered beta regression: a parsimonious, well-fitting model for continuous data with lower and upper bounds. Political Analysis, 31, 519–536.

Laurenson, K., Wood, M.J., Birkhead, T.R., Priestley, M.D.K., Sherley, R.B., Fayet, A.L., et al. (2025). Long-term multi-species demographic studies reveal divergent negative impacts of winter storms on seabird survival. Journal of Animal Ecology, 94, 139–153.

Layton-Matthews, K., Regan, C.E., Ballesteros, M., Hodges, K., Descamps, S., Anker-Nilssen, T., et al. (2025). Demographic responses of North Atlantic seabirds to seasonal ocean warming. Proceedings of the National Academy of Sciences, 122, e2507531122.

Leary, D.J. & Petchey, O.L. (2009). Testing a biological mechanism of the insurance hypothesis in experimental aquatic communities. Journal of Animal Ecology, 78, 1143–1151.

Lewis, S., Wanless, S., Wright, P., Harris, M., Bull, J. & Elston, D. (2001). Diet and breeding performance of black-legged kittiwakes Rissa tridactyla at a North Sea colony. Mar Ecol Prog Ser, 221, 277–284.

Lundberg, I., Johnson, R. & Stewart, B.M. (2021). What is your estimand? Defining the target quantity connects statistical evidence to theory. Am Sociol Rev, 86, 532–565.

Mallory, M.L., Robinson, S.A., Hebert, C.E. & Forbes, M.R. (2010). Seabirds as indicators of aquatic ecosystem conditions: A case for gathering multiple proxies of seabird health. Marine Pollution Bulletin, 60, 7–12.

Maunder, M.N. & Punt, A.E. (2004). Standardizing catch and effort data: a review of recent approaches. Fisheries Research, 70, 141–159.

Mckendrick, F.C., Descamps, S., Arnold, K.E., Jenouvrier, S., Harris, S.M., Bertrand, P., et al. (2026). Arctic-breeding black-legged kittiwakes show individual variation in foraging responses to glacial conditions without consequences for reproductive output. Oikos, 2026, e11558.

Mohan, K. & Pearl, J. (2014). Graphical models for recovering probabilistic and causal queries from missing data. In: Advances in neural information processing systems (eds. Ghahramani, Z., Welling, M., Cortes, C., Lawrence, N. & Weinberger, K.Q.). Curran Associates, Inc.

Mulder, J., Voelkle, M. & Hamaker, E.L. (2025). Time aggregation and missing time frames in causal research with panel data. PsyArXiv.

OSPAR. (2023). The OSPAR Regional Action Plan for Marine Birds in the North-East Atlantic (2024-2030).

Paleczny, M., Hammill, E., Karpouzi, V. & Pauly, D. (2015). Population trend of the world’s monitored seabirds, 1950-2010. PLOS ONE, 10, e0129342.

Pearl, J. (2009). Causality. 2nd edn. Cambridge University Press, Cambridge.

Ponchon, A., Grémillet, D., Christensen-Dalsgaard, S., Erikstad, K.E., Barrett, R.T., Reiertsen, T.K., et al. (2014). When things go wrong: intra-season dynamics of breeding failure in a seabird. Ecosphere, 5, art4.

R Core Team. (2025). R: A language and environment for statistical computing.

Reiertsen, T.K., Erikstad, K.E., Anker-Nilssen, T., Barrett, R.T., Boulinier, T., Frederiksen, M., et al. (2014). Prey density in non-breeding areas affects adult survival of black-legged kittiwakes Rissa tridactyla. Mar Ecol Prog Ser, 509, 289–302.

Reiertsen, T.K., Layton-Matthews, K., Erikstad, K.E., Hodges, K., Ballesteros, M., Anker-Nilssen, T., et al. (2021). Inter-population synchrony in adult survival and effects of climate and extreme weather in non-breeding areas of Atlantic puffins. Mar Ecol Prog Ser, 676, 219–231.

Rose, G.A. (2005a). Capelin (Mallotus villosus) distribution and climate: a sea “canary” for marine ecosystem change. ICES Journal of Marine Science, 62, 1524–1530.

Rose, G.A. (2005b). On distributional responses of North Atlantic fish to climate change. ICES Journal of Marine Science, 62, 1360–1374.

Sandvik, H., Erikstad, K., Barrett, R. & Yoccoz, N. (2005). The effect of climate on adult survival in five species of North Atlantic seabirds. Journal of Animal Ecology, 74, 817–831.

Sandvik, H., Erikstad, K.E. & Sæther, B.-E. (2012). Climate affects seabird population dynamics both via reproduction and adult survival. Mar Ecol Prog Ser, 454, 273–284.

Sandvik, H., Reiertsen, T.K., Erikstad, K.E., Anker-Nilssen, T., Barrett, R.T., Lorentsen, S.-H., et al. (2014). The decline of Norwegian kittiwake populations: modelling the role of ocean warming. Clim Res, 60, 91–102.

Scholz, M. & and Bürkner, P.-C. (2025). Prediction can be safely used as a proxy for explanation in causally consistent Bayesian generalized linear models. Journal of Statistical Computation and Simulation, 95, 1226–1249.

Searle, K.R., Regan, C.E., Perrow, M.R., Butler, A., Rindorf, A., Harris, M.P., et al. (2023). Effects of a fishery closure and prey abundance on seabird diet and breeding success: Implications for strategic fisheries management and seabird conservation. Biological Conservation, 281, 109990.

Simmonds, E.G., Adjei, K.P., Cretois, B., Dickel, L., González-Gil, R., Laverick, J.H., et al. (2024). Recommendations for quantitative uncertainty consideration in ecology and evolution. Trends in Ecology & Evolution, 39, 328–337.

Stan Development Team. (2025). Stan modeling language users guide and reference manual.

Sydeman, W.J., Schoeman, D.S., Thompson, S.A., Hoover, B.A., García-Reyes, M., Daunt, F., et al. (2021). Hemispheric asymmetry in ocean change and the productivity of ecosystem sentinels. Science, 372, 980–983.

Taylor, L.U. & Gnam, E. (2026). Prebreeding populations and the importance of life history for conserving imperiled seabirds. *Frontiers in Ecology and the Environment*, n/a, e70049.

Teplitsky, C., Mills, J.A., Alho, J.S., Yarrall, J.W. & Merilä, J. (2008). Bergmann’s rule and climate change revisited: Disentangling environmental and genetic responses in a wild bird population. Proceedings of the National Academy of Sciences, 105, 13492–13496.

Vaisey, S. & Miles, A. (2017). What you can—and can’t—do with three-wave panel data. Sociological Methods & Research, 46, 44–67.

Villejo, S.J., Martino, S., Illian, J., Ryan, W. & Lindgren, F. (2025). Validating uncertainty propagation approaches for two-stage Bayesian spatial models using simulation-based calibration. arXiv e-prints, arXiv:2502.18962.

Watts, K., Whytock, R.C., Park, K.J., Fuentes-Montemayor, E., Macgregor, N.A., Duffield, S., et al. (2020). Ecological time lags and the journey towards conservation success. Nature Ecology & Evolution, 4, 304–311.

Wood, M.J., Canonne, C., Besnard, A., Lachish, S., Fairhurst, S.M., Liedvogel, M., et al. (2021). Demographic profiles and environmental drivers of variation relate to individual breeding state in a long-lived trans-oceanic migratory seabird, the Manx shearwater. PLOS ONE, 16, e0260812.

Wooldridge, J.M. (1999). Distribution-free estimation of some nonlinear panel data models. Journal of Econometrics, 90, 77–97.

Yao, Y., Vehtari, A., Simpson, D. & Gelman, A. (2017). Using stacking to average Bayesian predictive distributions. arXiv e-prints, arXiv:1704.02030.

