## Supplementary Material for "Time-lagged causal inference corroborates presumed drivers of North Atlantic seabird ecology"

#### Contents

**S1: Causal justifications** 2

**S2: Supplementary methods** 6

**S3: Supplementary figures** 18

**S4: Supplementary tables** 31

**Supplementary references** 42

#### S1: Causal justifications

Our directed acyclic graph (DAG - Figure 1) was defined using a combination of established correlative evidence in the literature and natural history observations. This section describes and justifies each linkage. We should also clarify that as our measurements are limited to breeding adults, we assume all causal links are acting at the level of the adult population, unless explicitly stated. Variables present in the DAG are highlighted in bold.

**Environmental justifications:**

- **Climate** is a latent variable encompassing the entirety of the (assumed) physical environmental conditions. As climate is the long term average meteorology and weather (IPCC 2023), we represent climate as an unmeasured correlated error between temperature and weather (e.g. precipitation) variables. Climate is disambiguated between **breeding** and **non-breeding seasons** to reflect the fact that while birds are localised around the colony during the breeding season, over the non-breeding season they may travel large distances across the North Atlantic (Fayet et al. 2017). There is also a temporal component where non-breeding season conditions precede breeding season conditions, leading to potential endogeneity if not considered.
- **Prey fish accessibility** is the assumed primary driver of seabird declines (Cury et al. 2011; Reiertsen et al. 2014). However, it is challenging to accurately estimate fish availability given their wide spatio-temporal distribution (Maunder & Punt 2004), and the preference of Atlantic puffins (*Fratercula arctica*) and Black-legged kittiwakes (*Rissa tridactyla*) towards young fish (those <= 3 years old) typically unsampled in fishing data (Coulson 2011; Harris & Wanless 2011).
- **Zoo(phyto)plankton density** is known to drive fish abundance (Bachiller et al. 2016), which in turn is driven by **sea surface temperature** (**SST** - Broms & Melle 2007). Here, we focus on SST over plankton density primarily because of the broad correlative evidence for negative SST impacts on seabirds (i.e. abundance, breeding success, mass & survival decline - Fayet et al. 2021; Gibson et al. 2023; Irons et al. 2008; Sandvik et al. 2005; Sydeman

et al. 2021; Teplitsky et al. 2008), but assume that SST can have both direct and indirect impacts. For example, plankton density represents a fish food resource (Bachiller et al. 2016) which if disappears or moves, could lead to changing prey fish abundance and distribution (Rose 2005; Slotte et al. 2025). This is the indirect effect of SST on seabirds. Conversely, SST can have a direct effect on fish via thermal stress. This could manifest as mortality via physiological stress (Alfonso et al. 2021), or as a behavioural shift to deeper water (Freitas et al. 2021; Rose 2005).

- **Nutrients** are also impactful upon plankton density (Goldenberg et al. 2022). We assume North Atlantic nutrient concentration is primarily dependent on **upwellings** for their source (Pelegrí et al. 2006), and that concentration is dispersed by ocean currents/gyres (Johnson et al. 2013). We also believe ocean circulation directly influences fish locations (Cowen & Sponaugle 2009).
- Known *F. arctica* and *R. tridactyla* prey (*Ammodytes sp.*, *Clupea harengus*, and *Mallotus villosus*) are target species for **fishing** (Barrett & Krasnov 1996) which directly removes fish individuals from the system. This consequently impacts prey fish availability and quality (e.g. fish mass can shrink in response to overfishing - Engelhard & Heino 2004).
- **Air temperature** at the colony is a plausible physiological stress upon adult seabirds (Jones et al. 2018; Renner et al. 2024), especially as one parent remains at the nest while the other forages during the first part of chick-rearing (Coulson 2011; Harris & Wanless 2011). *Fractercula arctica* can probably minimise some of the impact via its burrows (Harris & Wanless 2011), but *R. tridactyla* have no such protection given they nest on cliff faces (Coulson 2011).
- **Precipitation** can also potentially drown chicks, wash away *R. tridactyla* nests or submerge *F. arctica* burrows. This will suppress breeding success.
- **Storms** during the non-breeding season are an emerging impact upon North Atlantic seabirds (Laurenson et al. 2025; Reiertsen et al. 2021). This impact is believed to act through both an indirect process (via prey fish availability) where fish are forced deeper and/or wave conditions are not conducive for seabird foraging, and directly as a physiological stress (birds exert extra energy to combat winds for example). Both these impacts affect adults and non-breeding juveniles but, as juveniles are not monitored, the outcome differs depending on the

bird's maturity. For example, abundance is influenced by juvenile mortality as juveniles do not contribute to the population measurement until they mature (Lorentsen & Christensen-Dalsgaard 2009), whereas overall survival is estimated solely from adults. We therefore assume survival is impacted by non-breeding season prey availability while breeding adult abundance is influenced by juvenile survival.

- During the breeding season, **wind speed** will also influence foraging decisions similarly to storms, with *R. tridactyla* choosing not to forage in adverse wind conditions (Collins et al. 2020).
- If **SST** and **air temperature** are expected to influence breeding birds, then it stands to reason that these variables are also influential in the non-breeding season.
- **Predation** (or disease) is known to alter both species' survival. In the North Atlantic, Great Skuas (*Catharacta skua*) are the primary predators (Oro & Furness 2002; Phillips et al. 1999). **Behaviour** also influences the likelihood of chick **predation** as adults with high foraging efforts attend their offspring less well.

**Intrinsic justifications:**

- We believe that intrinsic characteristics/measurements (behaviour, morphology, abundance etc.) influence one another, but because DAGs are 'acyclic' (they cannot have cyclical relationships), we are required to impose an order of changes. Fortunately, Cerini et al. (2023) and Stollewerk et al. (2025) suggest a predictable sequence of intrinsic changes is present.
- Specifically, they suggest **behaviour** at time t changes drive changes in **morphology** as behaviour is the most labile intrinsic characteristic available to a bird (Wong & Candolin 2015). This relationship has been demonstrated for seabirds (Fayet et al. 2016, 2021), where the energetic cost of foraging correlates with bird condition at a given time. As prey availability likely alters the foraging effort required to maintain body condition (Fayet et al. 2021), we have introduced the prey availability linkage via the foraging effort node.
- **Morphology** (body mass) then represents the allocation tradeoff between self maintenance and reproductive effort. A bird's body mass therefore influences both survival and breeding success (Kooijman et al. 2008), with different individuals and species potentially allocating

differentially.

- Finally, both **breeding success** and **survival** influence population **abundance**, but the lagged effect may differ given the recruitment effects in our focal seabird populations.

Endogeneity also exists given our belief that Cerini et al. (2023)'s assertions are true; historic intrinsic measurements can influence future intrinsic measurements. We assume that energy availability is the mediating node between time points, energy which we further assume is correlated with mass (Labocha & Hayes 2012). We therefore suggest that past energy (represented by body mass) influences current behaviour and body mass but not other intrinsic measurements as those are driven by the current body mass.

#### S2: Supplementary methods

##### Foraging effort

Bird location and behaviour was estimated using global location sensors (GLS), which record geographical position, light level and immersion data. Light and immersion data can be used to estimate behaviour, such as resting, travelling flight, and foraging (Christensen-Dalsgaard et al. 2018; Fayet et al. 2016). Foraging behaviour is thought to be influenced by exposure to extrinsic stressors (Chivers et al. 2012; Fayet et al. 2021), and foraging effort was therefore selected as the focal behavioural response. Each GLS report was assigned a behaviour based upon the combination of light level and conductivity detected by the immersion sensor. For puffins, following Fayet et al. (2016), a recording was classified as 'foraging' if light level was greater than 0 (i.e. there was light) and immersion was greater than 0.02 but less than 0.98 (i.e. the bird performed a sequence of short dry flights while searching for prey with short wet bouts of on/underwater activity). For kittiwakes, which are surface feeders and spend less time in the water than puffins (pursuit-divers) (Christensen-Dalsgaard et al. 2018; Fayet et al. 2016), the immersion thresholds were adjusted to greater than 0 but less than 1 so that any record including contact with both air and water was classified as foraging. We then calculated the number of daylight hours available to a bird per day using the suncalc v.0.5.1 (Thieurmel & Elmarhraoui 2022) R v.4.5.1 (R Core Team 2025) package and estimated the daily proportion of daylight hours each bird was foraging as a proxy for foraging effort. 341 puffin and 603 kittiwake individuals were tracked across all colonies.

We then used the geographical position recordings to estimate the utilization kernels for each population (Supplementary Material S4: Figure S1). As breeding puffins and kittiwakes are both central place foragers (Coulson 2011; Harris & Wanless 2011), flights during the breeding season are expected to be foraging trips, and thus location data over the sea is directly linked to their foraging behaviour. Raw positions were extracted and cleaned following Bråthen et al. (2021), with missing locations were interpolated based on the Informed Random Movement Algorithm (IRMA), which considers information on light levels, land masks and saltwater immersion data to infer colony presence and longitude during the equinoxes. Missing locations were replaced by plausible estimates using this approach (Fauchald et al. 2019). Utilization kernels were estimated via the utilization distribution provided by the adehabitatHR v.0.4.22 (Calenge 2024) R package. This kernel estimator represents a probability distribution which we cropped to the 50% quantile as the classic definition of 'core area'. Focus on this core area was also necessitated by the low resolution (~158km) of the GLS loggers used (Fayet et al. 2016). We did not assume time varying kernels given the low sample sizes of certain populations in certain years. Consequently, only a single breeding season kernel was estimated for each population, pooling individual tracks across all years. We repeated this process for the non-breeding season.

**Defining the breeding season**

Breeding and non-breeding season dates were classified following Fauchald et al. (2019), using the 5-day rolling mean across individual birds' GLS conductivity timeseries in combination with the Lavielle partitioning algorithm (Barraquand & Benhamou 2008). In brief, this process classifies periods of mostly wet and periods of mostly dry, with the dry periods assumed to represent colony attendance and therefore the breeding season. All seabird measurements were classified using these date ranges into breeding and non-breeding periods.

**Morphology**

For morphometrics, breeding adults were caught using nest traps during the incubation period (Lorentsen & Christensen-Dalsgaard 2009). Once caught, birds were weighed, and their head and bill length measured. A total of 9960 measurements were taken across 2989 puffin individuals and 3733 kittiwake individuals. Sexing was performed using blood derived DNA tests (Anker-Nilssen et al. 2018) with ~53 % of sampled birds sexed.

**Breeding success**

Breeding success was recorded in sampling plots, based on a sample of nests that are monitored at multiple instances over the season. For kittiwakes, breeding success was measured as either the number of medium or large kittiwake chicks, or chicks older than 15 days, per nest (Walsh et al. 1995). Puffin breeding success was measured as either the number of chicks 20/30 days or older, medium/large-sized chicks per egg laid, the number of chicks (Ponchon et al. 2014). While kittiwakes can lay up to three eggs, most only lay up to two (Coulson 2011). Therefore, kittiwakes breeding success can result in breeding success values larger than one, which we divided by two to standardise.

##### Survival

Survival was tracked via individual resighting histories using unique numbered metal rings and individually coded colour rings readable through telescopes or binoculars. Visual searches were made for marked birds each breeding season, and observations recorded. In total, 2726 puffin and 3579 kittiwake individuals were tracked across all colonies.

##### Population size

We summed apparently occupied nests across all plots within a colony to give a representation of minimum bird population size - i.e. the maximum number of observed pairs is the minimum present in the colony. As the number of plots can vary between colonies and years, we retained information on sampling effort to control for changing effort in downstream analyses.

Røst kittiwakes required additional data processing as colony-level breeding success and population size was measured across two sites (Vedøy and Harbour) which display different trends and consequently different insights into the fate of this colony (i.e. Vedøy supported a large population which declined over 40 years to extinction in 2020, whereas Harbour is a much smaller colony that emerged in 1980, grew, and now is in decline). While maximum apparently occupied nests were similar between the two sites, Vedøy counts represented 2% sampling effort (estimated from a true versus sampling plot count comparison made in 1988) whereas it was 100% at Harbour. We therefore estimated Røst kittiwake population size as the weighted average between the two sites using the following equation:

$$x_{røst} = \frac{w_{vedøy} \cdot x_{vedøy} + w_{harbour} \cdot x_{harbour}}{w_{vedøy} + w_{harbour}}$$

$$w_{vedøy} = \frac{\Lambda \cdot nplots_{vedøy}}{\Lambda \cdot nplots_{vedøy} + nplots_{harbour}}$$

$$w_{harbour} = \frac{nplots_{harbour}}{\Lambda \cdot nplots_{vedøy} + nplots_{harbour}}$$

$$\Lambda = \frac{x_{true}}{x_{sample}}$$

where $w$ is the site's weight, $x$ is the site's population size summed across all sampling plots, *nplots* is the number of sampling plots at the site, and $\Lambda$ is the ratio of true kittiwake count to total sample plot count on Vedøy in 1988.

Røst kittiwake breeding success was then estimated as a weighted average relative to each site's time varying contribution to the colony using the equation:

$$bs_{røst,i} = \frac{\gamma_{vedøy,i} \cdot bs_{vedøy,i} + \gamma_{harbour,i} \cdot bs_{harbour,i}}{\gamma_{vedøy,i} + \gamma_{harbour,i}}$$

$$\gamma_{vedøy,i} = \frac{w_{vedøy} \cdot x_{vedøy,i}}{x_{røst,i}}$$

$$\gamma_{harbour,i} = \frac{w_{harbour} \cdot x_{harbour,i}}{x_{røst,i}}$$

where $w$ is the site weight, $x_i$ is the site's population size summed across all sampling plots in year $i$, $bs$ is the site's breeding success, and $\gamma_i$ is the site's relative contribution to the total Røst colony population size in year $i$.

##### Average population body mass

Average yearly population body mass was modelled as a multivariate normal mixture between bird body mass (g) and skull + bill length (mm) using the equation:

$$y_i \sim MVN(\mu_i, \Sigma) \; if \; sex_i \in \{0,1\}$$
$$\mu_i = (\mu_{mass[i]}, \mu_{size[i]})$$
$$\mu_{mass[i]} = \beta_{mass} + \beta_{yearM[i]} + \beta_{dayM[i]} + \beta_{indM[i]} + \beta_{sexM} \cdot sex_i$$
$$\mu_{size[i]} = \beta_{size} + \beta_{indS[i]} + \beta_{sexS} \cdot sex_i$$

$$y_i \sim MVN(\mu_{i,sex}, \Sigma) \cdot \lambda \; if \; sex_i \notin \{0,1\}$$
$$\mu_{i,sex} = (\mu_{mass[i],sex}, \mu_{size[i],sex})$$
$$\mu_{mass[i],sex=1} = \beta_{mass} + \beta_{yearM[i]} + \beta_{dayM[i]} + \beta_{indM[i]} + \beta_{sexM}$$
$$\mu_{mass[i],sex=0} = \beta_{mass} + \beta_{yearM[i]} + \beta_{dayM[i]} + \beta_{indM[i]}$$
$$\mu_{size[i],sex=1} = \beta_{size} + \beta_{indS[i]} + \beta_{sexS}$$
$$\mu_{size[i],sex=0} = \beta_{size} + \beta_{indS[i]}$$

$$sex_i \sim Bernoulli(\lambda)$$
$$\Sigma = \begin{bmatrix} \tau_{mass}^2 & \rho \cdot \tau_{mass} \cdot \tau_{size} \\ \rho \cdot \tau_{mass} \cdot \tau_{size} & \tau_{size}^2 \end{bmatrix}$$
$$\beta_{mass} \sim Normal(400, 100)$$
$$\beta_{yearM} \sim Normal(0, \sigma_{yearM})$$
$$\beta_{dayM} \sim Normal(0, \sigma_{dayM})$$
$$\beta_{indM} \sim Normal(0, \sigma_{indM})$$
$$\beta_{sexM} \sim Normal(0, 10)$$
$$\beta_{size} \sim Normal(80, 25)$$
$$\beta_{indS} \sim Normal(0, \sigma_{indS})$$
$$\beta_{sexS} \sim Normal(0, 10)$$
$$\lambda \sim Uniform(0, 1)$$
$$\sigma \sim Exponential(1)$$
$$\tau_{mass} \sim Normal_+(25, 10)$$
$$\tau_{size} \sim Normal_+(1, 5)$$
$$\rho \sim LKJcorr(1)$$

where $y$ is a vector containing skull length and body mass at index $i$, $\mu$ is the location parameter, $\Sigma$ is the covariance matrix between skull length and body mass, and $\beta$ are linear predictor coefficients. $\tau$ is the scale parameter. $\rho$ is the residual correlation between the response variables, and $\lambda$ is the mixture probability which is equivalent to the sex ratio. $\sigma$ is the random effect variance.

**Average population foraging behaviour**

Average yearly foraging behaviour was modelled as a ordinal beta mixture using the equation:

$$y_i \sim OrderedBeta(\alpha, \gamma, \delta, \mu_i, \phi) \; if \; sex_i \in \{0, 1\}$$
$$\mu_i = \beta_0 + \beta_{year[i]} + \beta_{ind[i]} + \beta_{sex} \cdot sex_i$$

$$y_i \sim OrderedBeta(\alpha, \gamma, \delta, \mu_{i,sex}, \phi) \cdot \lambda \; if \; sex_i \notin \{0, 1\}$$
$$\mu_{i,sex=1} = \beta_0 + \beta_{year[i]} + \beta_{ind[i]} + \beta_{sex}$$
$$\mu_{i,sex=0} = \beta_0 + \beta_{year[i]} + \beta_{ind[i]}$$

$$\alpha = 1 \cdot (\mu_i - k_1) \; if \; y_i = 0$$
$$\delta = ((\mu_i - k_1) - (\mu_i - k_2)) \cdot Beta(\mu_i, \phi) \; if \; y_i \in (0, 1)$$
$$\gamma = 1 - (\mu_i - k_2) \; if \; y_i = 1$$

$$sex_i \sim Bernoulli(\lambda)$$

$$\beta_0 \sim Normal(0, 5)$$
$$\beta_{sex} \sim Normal(0, 10)$$
$$\beta_{year} \sim Normal(0, \sigma_{year})$$
$$\beta_{ind} \sim Normal(0, \sigma_{ind})$$
$$\lambda \sim Uniform(0, 1)$$
$$\phi \sim Exponential(1)$$
$$k_{1,2,3} \sim InducedDirichlet(1, 1, 1)$$
$$\sigma \sim Exponential(0.1)$$

where $y$ is proportion of daylight hours spent foraging at index $i$, $\alpha$ is the probability of an observation being 0, $\gamma$ is the probability an observation is the continuous range, $\delta$ is the probability an observation is 1, and $k$ is the ordered cutpoints between the components. $\mu$ is the location parameter with $\beta$ being the linear predictor coefficients, $\lambda$ the mixture probability which is equivalent to the sex ratio, $\phi$ is the precision parameter of the reparameterized *Beta* distribution, and $\sigma$ the random effect variance.

##### Average population survival

Average yearly survival was estimated from capture-mark-resight data using a Cormack-Jolly-Seber model that incorporated first-order Markovian trap dependence (Pradel & Sanz-Aguilar 2012) and transience effects (Pradel et al. 1997). Specifically, recapture probability at year t depended on whether an individual had been encountered at year t-1 using the following equation:

$$y_{i,t} \sim Bernoulli(p_{i,t-1})$$

$$1 \sim Bernoulli(\phi_{i,t-1})$$

$$1 \sim Bernoulli(\chi_i)$$

$$p_{i,t} = invlogit(\beta_{dependency[i]} + \sigma_{p,t})$$

$$\phi_{i,t} = invlogit(\mu_{transience[i]} + \sigma_{\phi,t})$$

$$\chi_{i,t} = (1 - \phi_{i,t}) + \phi_{i,t} \cdot (1 - p_{i,t}) \cdot \chi_{i,t+1}$$

$$\beta_{dependency} \sim Normal(0, 1)$$

$$\mu_{transience} \sim Normal(0, 1)$$

$$\sigma_p \sim Normal_+(5, 3)$$

$$\sigma_\phi \sim Normal_+(5, 3)$$

where $y$ is the capture history of a bird $i$ at timepoint $t$. $p$ is consequently the encounter probability, $\phi$ is survival probability, and $\chi$ is the probability of not capturing an individual. $\beta$ is the average trap dependence, $\mu$ is the average survival while $\sigma$ is the variance across years.

##### Gaussian state space interpolation

Gaussian distributed measurements (i.e. body mass) were interpolated using the following equation:

$$y_t \sim Normal(LV_t, \sigma_y)$$
$$LV_1 \sim Normal(y_1, 1)$$
$$LV_{2:t} = LV_{t-1} + \mu + \epsilon_t$$

$$\mu \sim Normal(0, 1)$$
$$\epsilon \sim Normal(0, \sigma_x)$$
$$\sigma_{x,y} \sim InvGamma_+(6, 6)$$

where $y$ is the observed measurement at time $t$, $LV$ is the latent process, $\mu$ is the process trend and $\epsilon$ is the error around the latent process for each time point. $\sigma_x$ is the standard deviation of that process error whereas $\sigma_y$ is the standard deviation of the observation error.

##### Poisson state space interpolation

Count measurements (i.e. population size) were interpolated using the following equation:

$$y_t \sim Poisson(exp(LV_t))$$
$$LV_1 \sim Normal(0, 1)$$
$$LV_{2:t} = LV_{t-1} + \mu + \epsilon_t$$

$$\mu \sim Normal(0, 1)$$
$$\epsilon \sim Normal(0, \sigma_x)$$
$$\sigma_x \sim Normal_+(0, 1)$$

where $y$ is the observed measurement at time $t$, $LV$ is the latent process, $\mu$ is the process trend and $\epsilon$ is the error around the latent process for each time point. $\sigma_x$ is the standard deviation of that process error.

##### Ordered beta state space interpolation

Proportional measurements (i.e. foraging effort, breeding success and survival) were interpolated using the following equation:

$$
\begin{gathered}
y_t \sim OrderedBeta(\alpha, \gamma, \delta, LV_t, \phi) \\
LV_1 \sim Normal(y_1, 1) \\
LV_{2:t} = LV_{t-1} + \mu + \epsilon_t \\
\alpha = 1 \cdot (LV_t - k_1) \; if \; y_t = 0 \\
\delta = ((LV_t - k_1) - (LV_t - k_2)) \cdot Beta(LV_t, \phi) \; if \; y_t \in (0, 1) \\
\gamma = 1 - (LV_t - k_2) \; if \; y_t = 1 \\
\\
\mu \sim Normal(0, 1) \\
\phi \sim Exponential(0.1) \\
k_{1,2,3} \sim InducedDirichlet(1, 1, 1) \\
\epsilon \sim Normal(0, \sigma_x) \\
\sigma_x \sim InvGamma_{+}(6, 6)
\end{gathered}
$$

where $y$ is the observed measurement at time $t$, $\alpha$ is the probability of an observation being 0, $\gamma$ is the probability an observation is the continuous range, $\delta$ is the probability an observation is 1, and $k$ is the ordered cutpoints between the components. $LV$ is therefore the latent process, $\mu$ is the process trend and $\epsilon$ is the error around the latent process for each time point. $\sigma_x$ is the standard deviation of that process error and $\phi$ is the precision parameter of the reparameterized *Beta* observation distribution.

##### Posterior fusion

To propogate error from the sequence of models fit, we used the capability of Bayesian modelling to pass entire posteriors between modelling stages. However, due to the state dependent nature of state-space modelling, we were forced to use a two step modelling process rather than the preferred joint-modelling method (Simmonds et al. 2024). Each draw from the population average models (Section *Obtaining population level estimates from individual level data*) was considered a time series and then interpolated using a separate state-space model (Section *Interpolation of missing years*).

This creates a distribution of distributions which can then be combined into a single posterior distribution via posterior fusion (Villejo et al. 2025). This is possible as each distribution is simply a different realisation of a shared process. A graphical example of this methodology is provided in Supplementary Material S4: Figure S3.

##### Fixed effect panel models

Panel models were fit separately for puffins and kittiwakes using the following model skeleton:

$$y_i \sim \beta_0 + \delta_{population} + \beta_1 \cdot x_i + \beta_W \cdot W_i$$

that varied depending on the distribution family and minimum adjustment set (see Supplementary Material S2 for exact model details). $y$ is the response for observation $i$, $x$ is the hypothesised causal variable and $W$ is a matrix containing the time varying adjustment set. Consequently, $\delta_{population}$ is the colony unit level fixed effect, $\beta_1$ is the causal effect for $x$, and $\beta_W$ are a varying number of covariate coefficients depending on the number of variables in the adjustment set $W$. For example, assuming our DAG's accuracy, to estimate the influence of SST on foraging effort, wind speed should be adjusted as there is a backdoor path via the unmeasured climate (Figure 1, Supplementary Material S3: Figure S7A). Conversely, to estimate the effect of SST upon population count (Supplementary Material S3: Figure S7B,C), we included colony air temperature, wind speed, and precipitation. Adjustment sets for fishery pressure and storm duration can be found in Supplementary Material S3: Figure S8-S9.

Gaussian distributions were fit to body mass data (Dee et al. 2023), log Gaussian to count data (Wooldridge 1999), and ordered beta distributions to foraging effort, breeding success and survival data (Kubinec 2023).

##### Count panel model

One way fixed effect panel models were fit to logged counts using the equation:

$$log(y_i) \sim Normal(\mu_i, \sigma)$$

$$\mu_i = \beta_0 + \delta_{population} + \beta_1 \cdot x_i + \beta_C \cdot X_i + offset\left(log\left(nplots\right)\right)$$

$$\beta_0 \sim Normal(0, 3)$$
$$\delta_{population} \sim Normal(0, 3)$$
$$\beta_{1,C} \sim Normal(0, 3)$$
$$\sigma \sim Exponential(1)$$

where $y$ is the observed measurement at index $i$, $x_i$ is the causal variable of interest and $X$ is a matrix of covariates. $\beta_0$ is therefore the intercept,$\delta_{population}$ is the population fixed effect, $\beta_1$ is the causal coefficient of interest and $\beta_C$ are coefficients for the covariates. $\sigma$ is the variance of the error distribution.

**Gaussian panel model**

One way fixed effect panel models were fit to average body mass (g) using the equation:

$$y_i \sim Normal(\mu_i, \sigma)$$
$$\mu_i = \beta_0 + \delta_{population} + \beta_1 \cdot x_i + \beta_C \cdot X_i$$
$$\beta_0 \sim Normal(400, 25)$$
$$\delta_{population} \sim Normal(0, 50)$$
$$\beta_{1,C} \sim Normal(0, 50)$$
$$\sigma \sim Exponential(0.1)$$

where $y$ is the observed measurement at index $i$, $x_i$ is the causal variable of interest and $X$ is a matrix of covariates. $\beta_0$ is therefore the intercept, $\delta_{population}$ is the population fixed effect, $\beta_1$ is the causal coefficient of interest and $\beta_C$ are coefficients for the covariates. $\sigma$ is the variance of the error distribution.

**Proportion panel model**

One way fixed effect panel models were fit to average survival, breeding success and foraging effort using the equation:

$$y_i \sim OrderedBeta(\alpha, \gamma, \delta, \mu_i, \phi)$$

$$\mu_i = \beta_0 + \delta_{population} + \beta_1 \cdot x_i + \beta_C \cdot X_i$$
$$\alpha = 1 \cdot (\mu_i - k_1) \ if \ y_i = 0$$
$$\delta = ((\mu_i - k_1) - (\mu_i - k_2)) \cdot Beta(\mu_i, \phi) \ if \ y_i \in (0, 1)$$
$$\gamma = 1 - (\mu_i - k_2) \ if \ y_i = 1$$

$$\beta_0 \sim Normal(0, 5)$$
$$\delta_{population} \sim Normal(0, 5)$$
$$\beta_{1,C} \sim Normal(0, 5)$$
$$\phi \sim Exponential(0.1)$$
$$k_{1,2,3} \sim InducedDirichlet(1, 1, 1)$$

where $y$ is the observed measurement at index $i$, $x_i$ is the causal variable of interest and $X$ is a matrix of covariates. $\alpha$ is the probability of an observation being 0, $\gamma$ is the probability an observation is the continuous range, $\delta$ is the probability an observation is 1, and $k$ is the ordered cutpoints between the components.$\beta_0$ is therefore the intercept, $\delta_{population}$ is the population fixed effect, $\beta_1$ is the causal coefficient of interest and $\beta_C$ are coefficients for the covariates. $\phi$ is the precision parameter of the reparameterized $Beta$ observation distribution.

#### S3: Supplementary figures

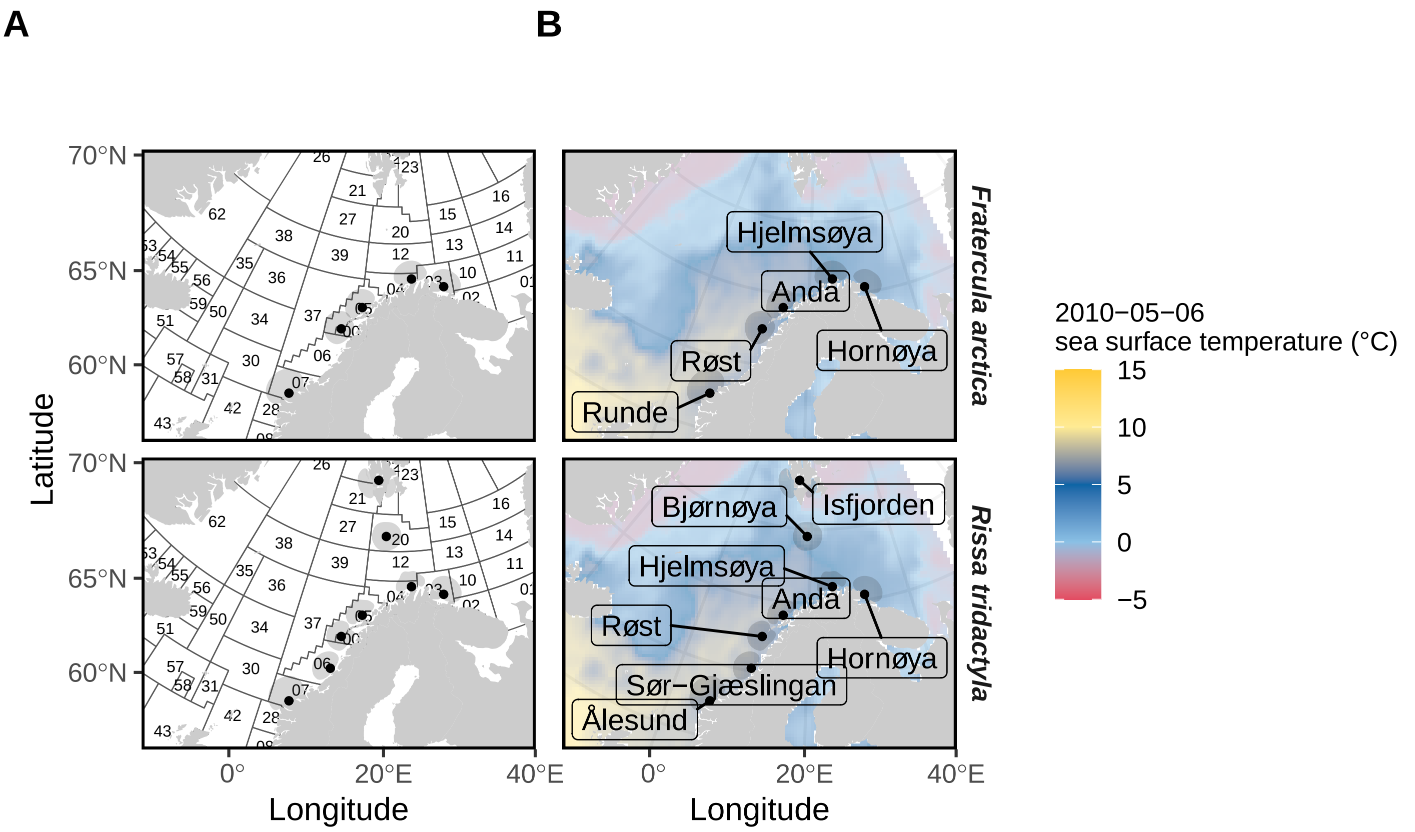


Figure S1: Colony locations and breeding season core-area kernels relative to A) Fiskeridirektoratet main fishing areas and B) mean Optimum Interpolation Sea Surface Temperature (°C) on 6th May 2010.

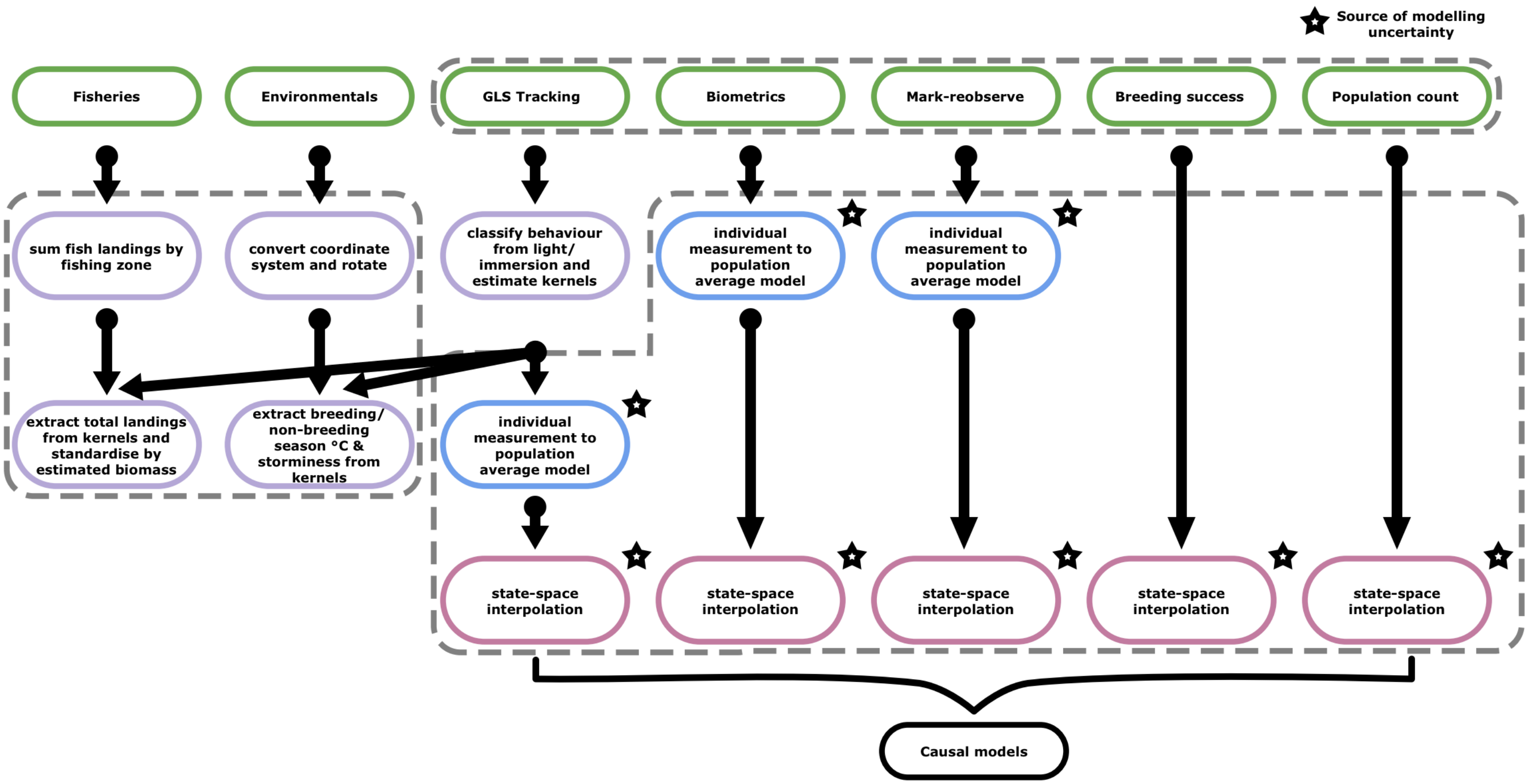


Figure S2: Visualisation of the analytical workflow. Raw data is either averaged using generalised linear mixed effect models prior to interpolation, or is interpolated directly. These interpolated time series are then analysed to identify the causal relationships.

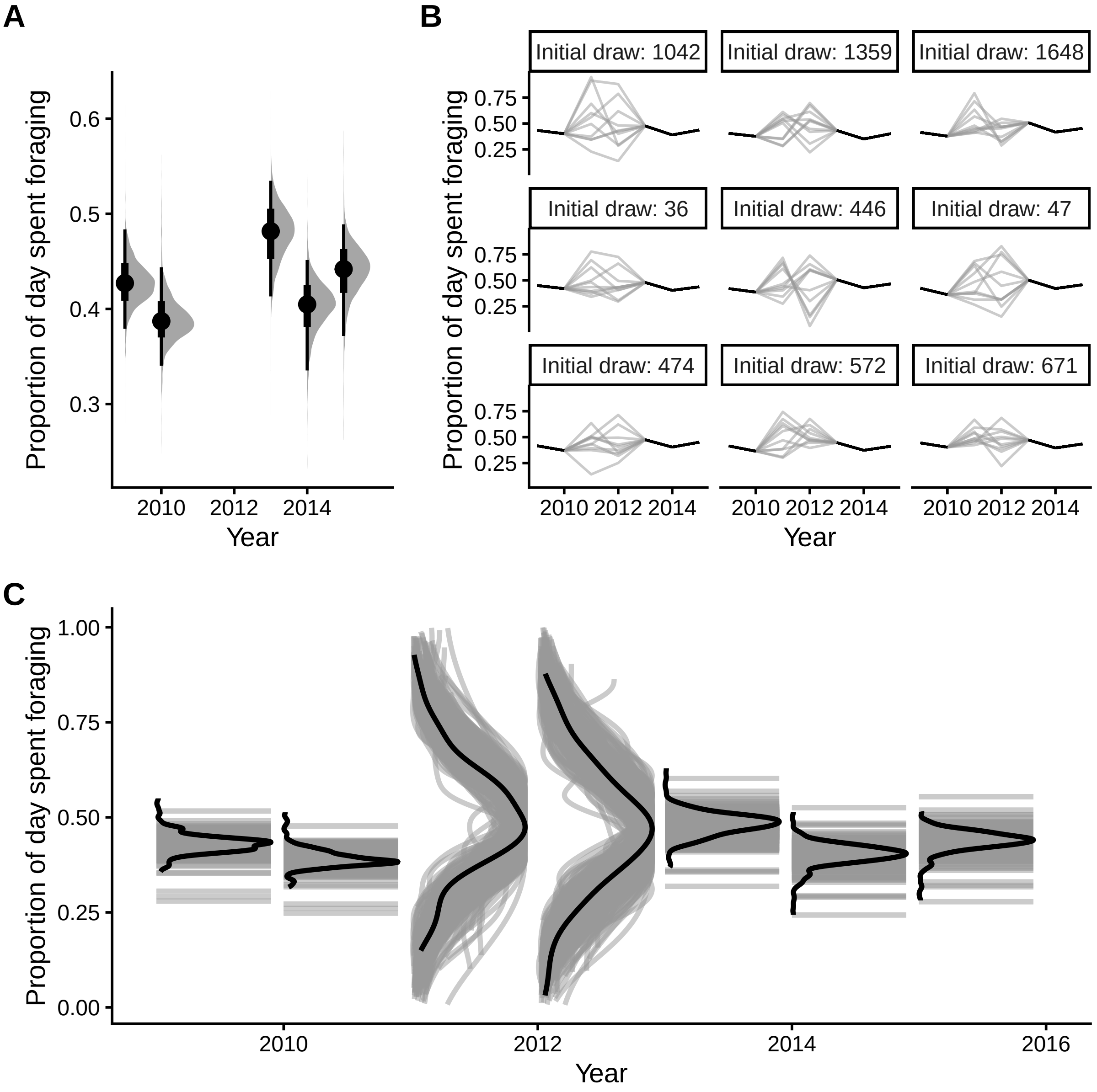


Figure S3: Example of error propagation across initial averaging models and state-space interpolation via posterior fusion. The behaviour estimates of *Rissa tridactyla* from Hjelmsøya are presented as an example. A) Output of the initial averaging model (Supplementary Material S1: Equation 3). Estimates are presented as density plots of model posterior distributions where points are median estimates, and error bars the 95% credible interval. Years 2011 and 2012 have no data and so are to be interpolated. B) As an example, nine random draws are taken from the distributions in panel A, to form nine time series (black lines) the model believes likely. The missing years are interpolated using a ordered beta state-space model (see Supplementary Material S1: Equation 6 for each draw time series, which results in a new distribution for each year. The state-space interpolations are presented as grey lines while the uninterpolated data are in black. C) The set of posterior distributions for each time point can be combined and resampled to form a 'fused posterior'. Light grey density plots are the posterior for each unique draw of the state space interpolation, while black density plots are the fused yearly posteriors.

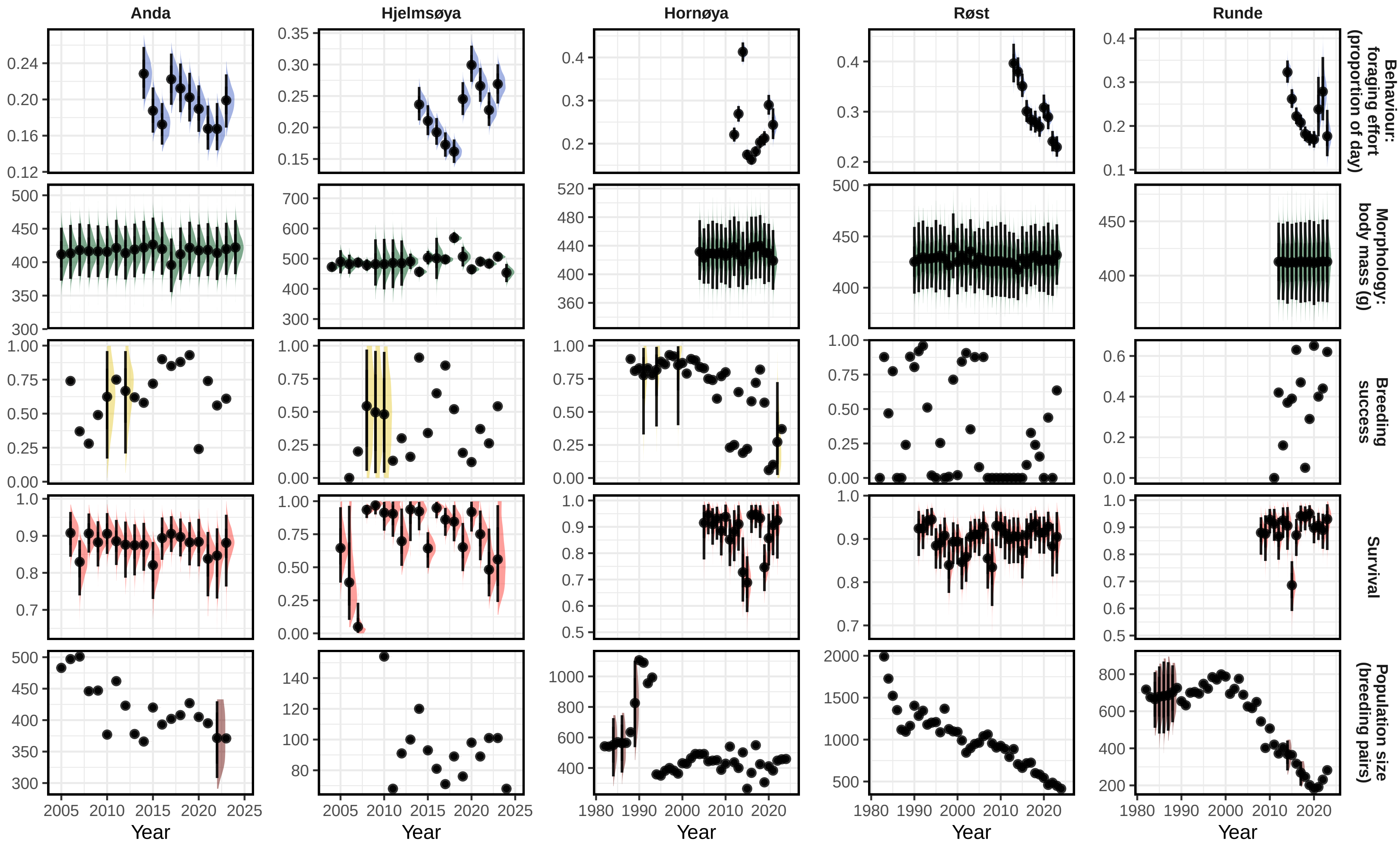


Figure S4: Estimated *Fratercula arctica* measurement time series following averaging and interpolation. Estimates are presented as density plots of model posterior distributions where points are median estimates, and error bars the 95% credible interval.

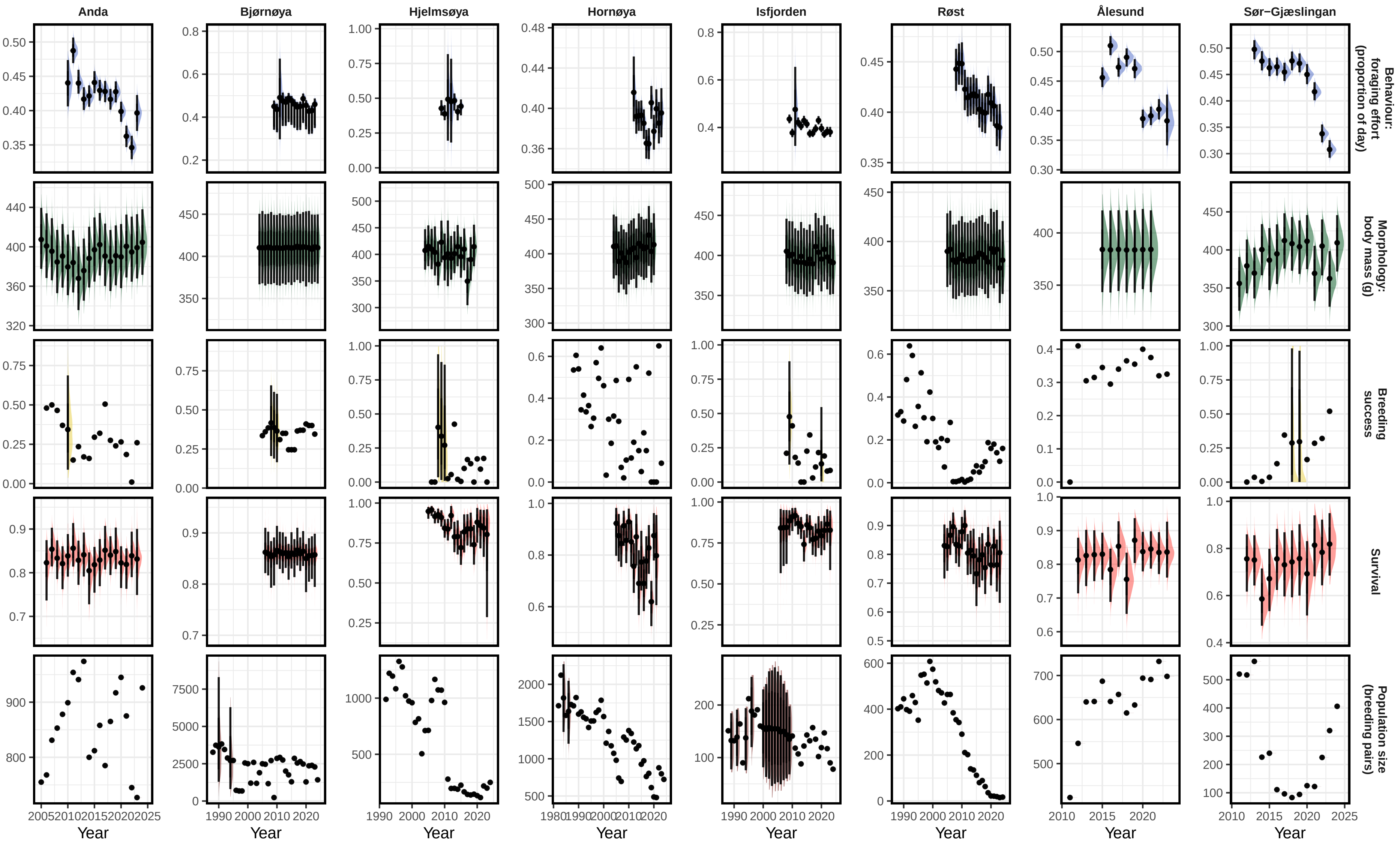


Figure S5: Estimated *Rissa tridactyla* measurement time series following averaging and interpolation. Estimates are presented as density plots of model posterior distributions where points are median estimates, and error bars the 95% credible interval.

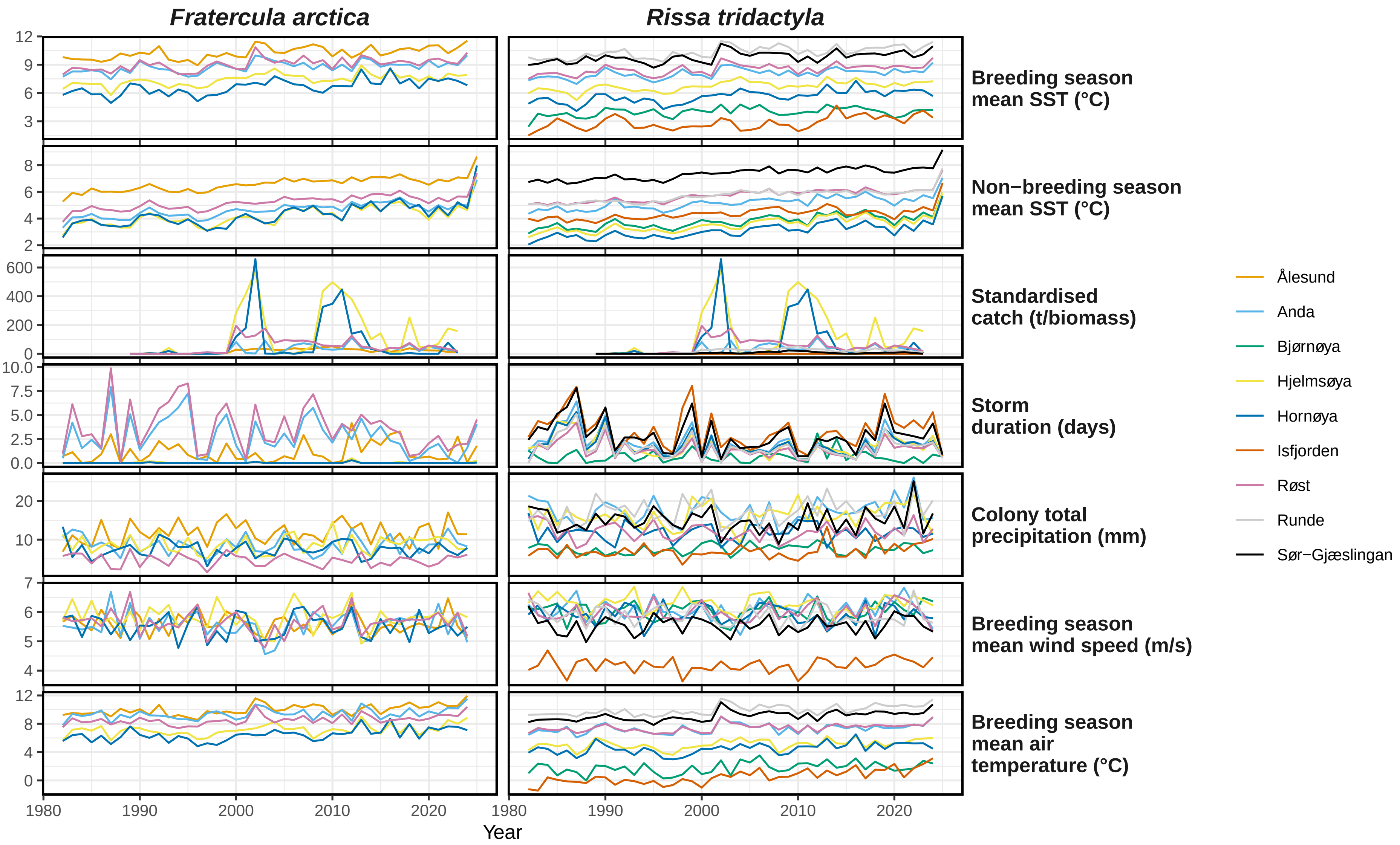


Figure S6: Stress trends across species and populations.

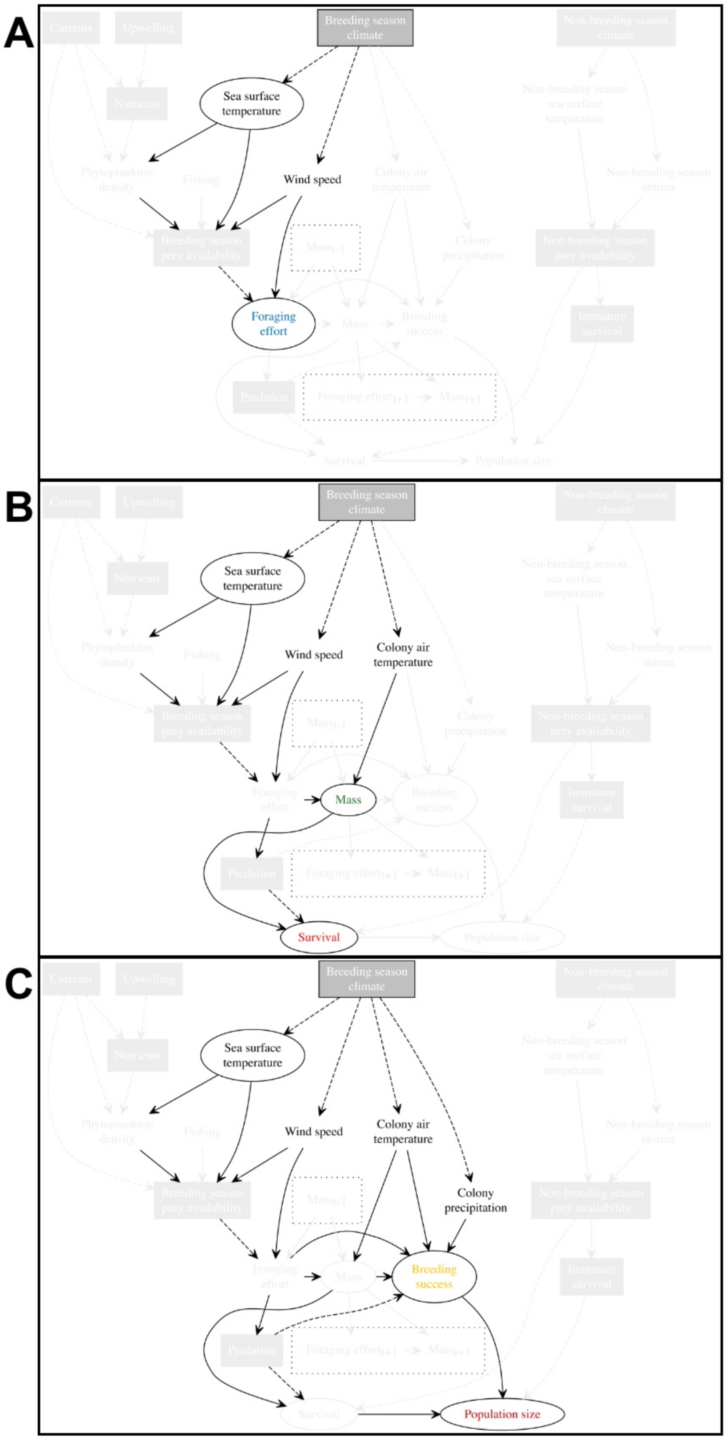


Figure S7: Selection of adjustment sets for sea surface temperature impacts implied by the DAG in Figure 1. The focal causal nodes are indicated by circles, unmeasured variables with grey boxes, and inappropriate covariates transparent. Required covariates are the remaining nodes. A) The adjustment set for behaviour (foraging effort). B) The adjustment set for morphology (body mass), and survival. C) The adjustment set for breeding success and abundance.

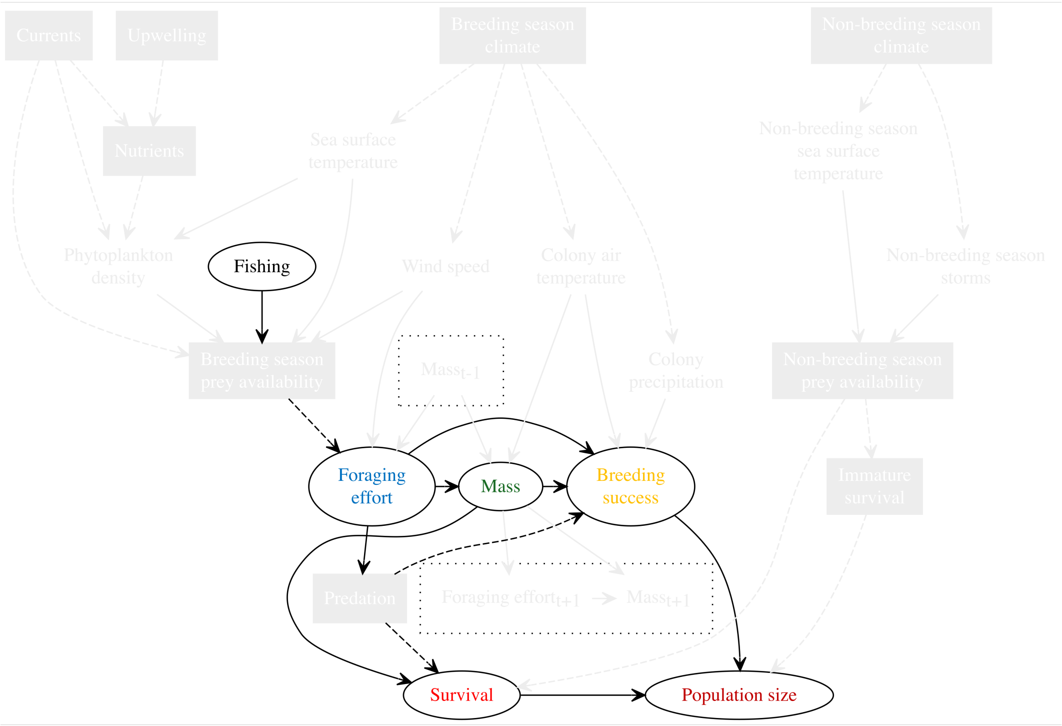


Figure S8: Selection of covariates for fishing pressure impacts implied by the DAG in Figure 1. The focal causal nodes are indicated by circles, unmeasured variables with grey boxes, and inappropriate covariates transparent. Required covariates are the remaining nodes.

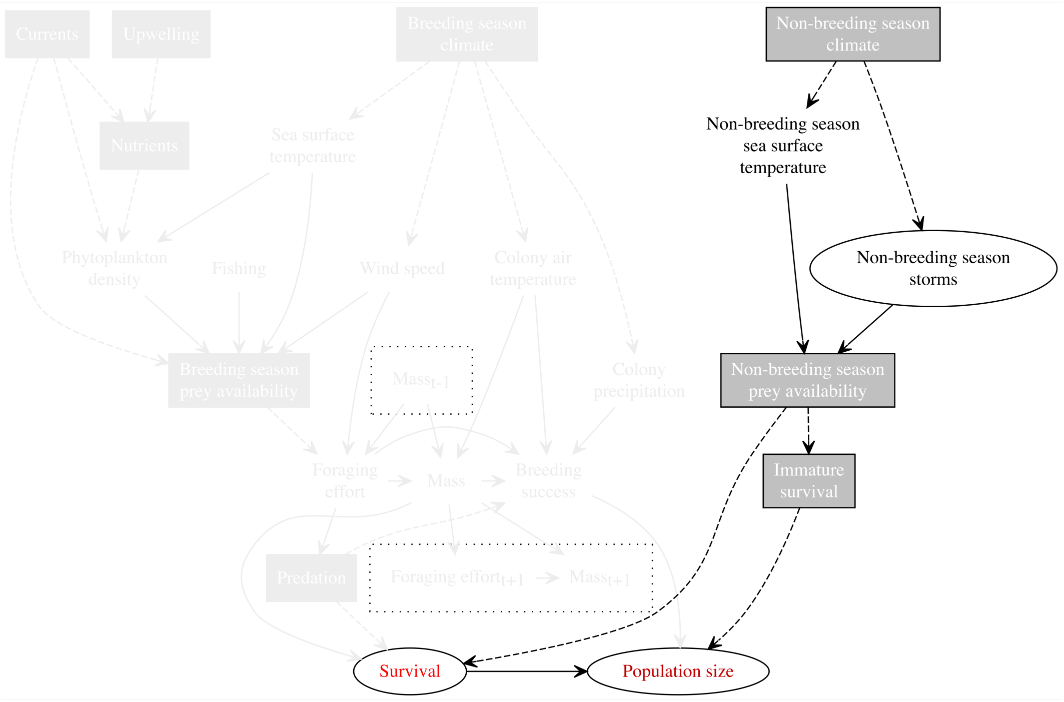


Figure S9: Selection of covariates for storm impacts implied by the DAG in Figure 1. The focal causal nodes are indicated by circles, unmeasured variables with grey boxes, and inappropriate covariates transparent. Required covariates are the remaining nodes.

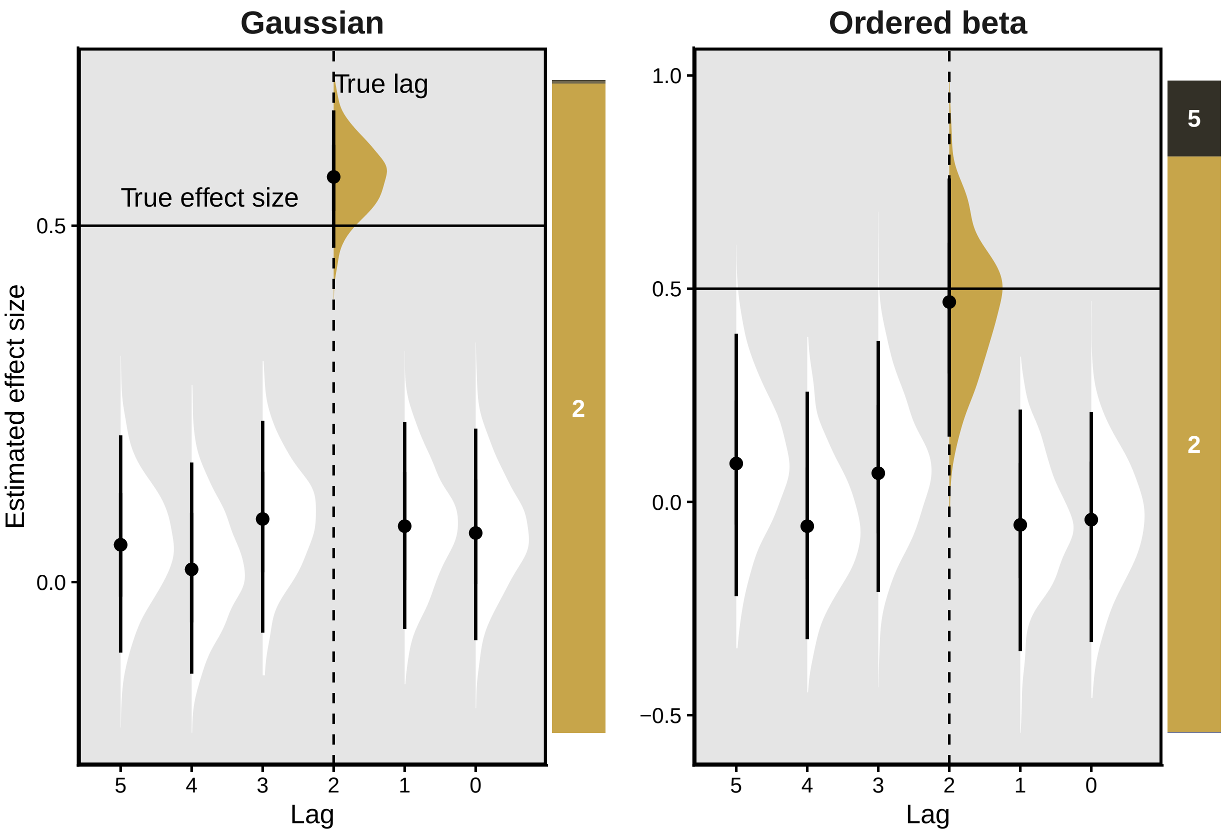


Figure S10: Demonstration of lagged causal model ability to correctly detect both the true causal effect and lag in simulated data. Generating equations: $\mu = 0.5 \cdot x_{t-2}$; $y \sim Normal(\mu, \sigma = 0.5)$; $y \sim OrderedBeta(\mu, \phi = 0.5)$. Six models are fit to each dataset (one for each lag), with each model assigned a weight based upon its leave-one-out predictive ability. Estimates are presented as density plots of model posterior distributions where points are median estimates, and error bars the 95% credible interval. Stacked bar charts are the model weights, labelled by each lag. These simulation experiments demonstrate that the correct causal effect is estimated (i.e 0.5), and the correct lag (i.e. $t - 2$) is the dominant contributor (i.e. fills the majority of the stacked bar).

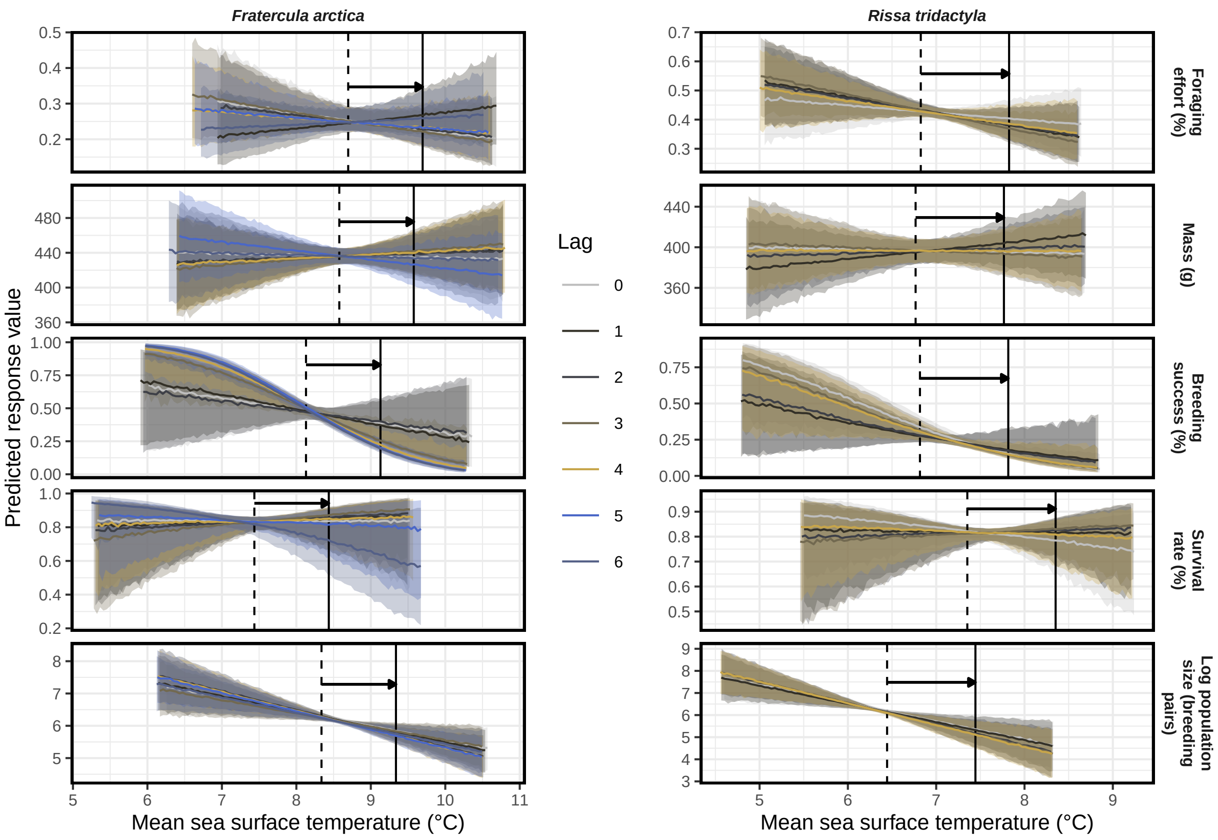


Figure S11: Marginal curves across the range of sea surface temperatures observed. Arrows indicate the predictions that inform the average treatment effect.

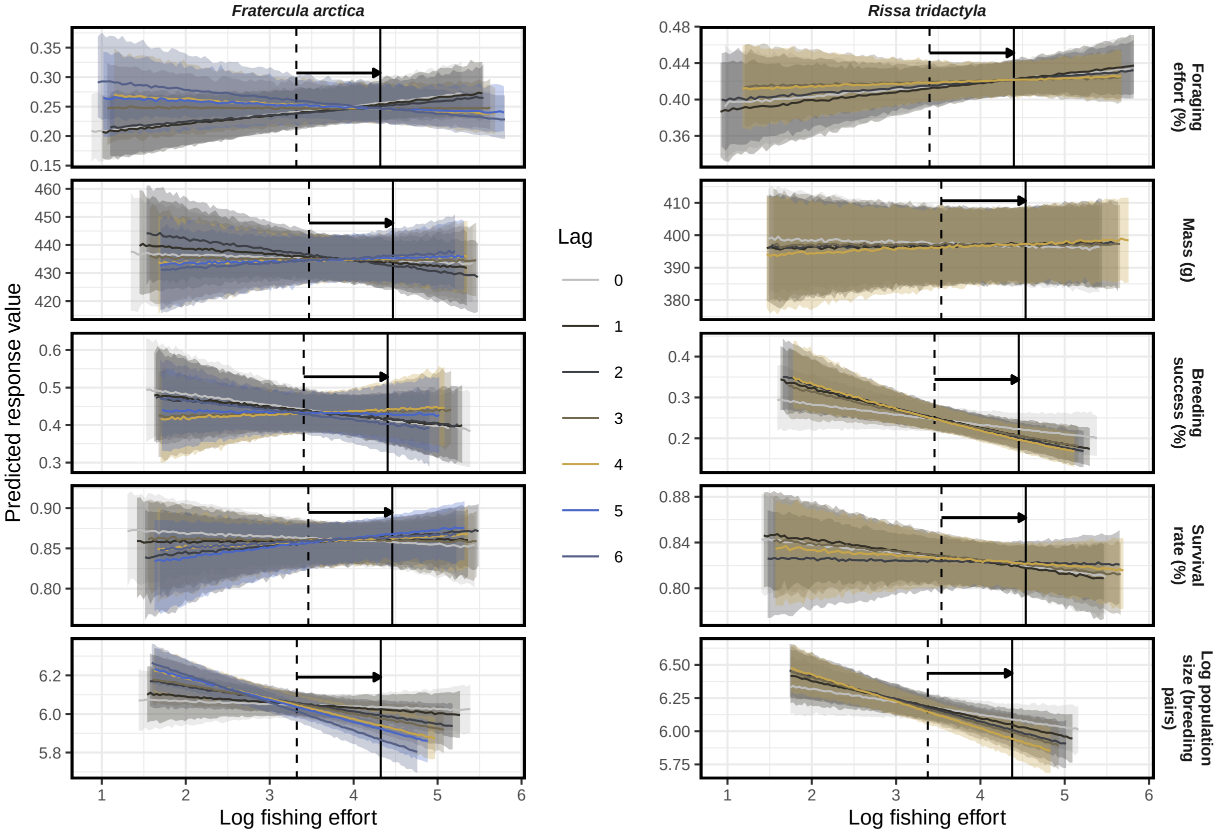


Figure S12: Marginal curves across the range of fishing efforts observed. Arrows indicate the predictions that inform the average treatment effect.

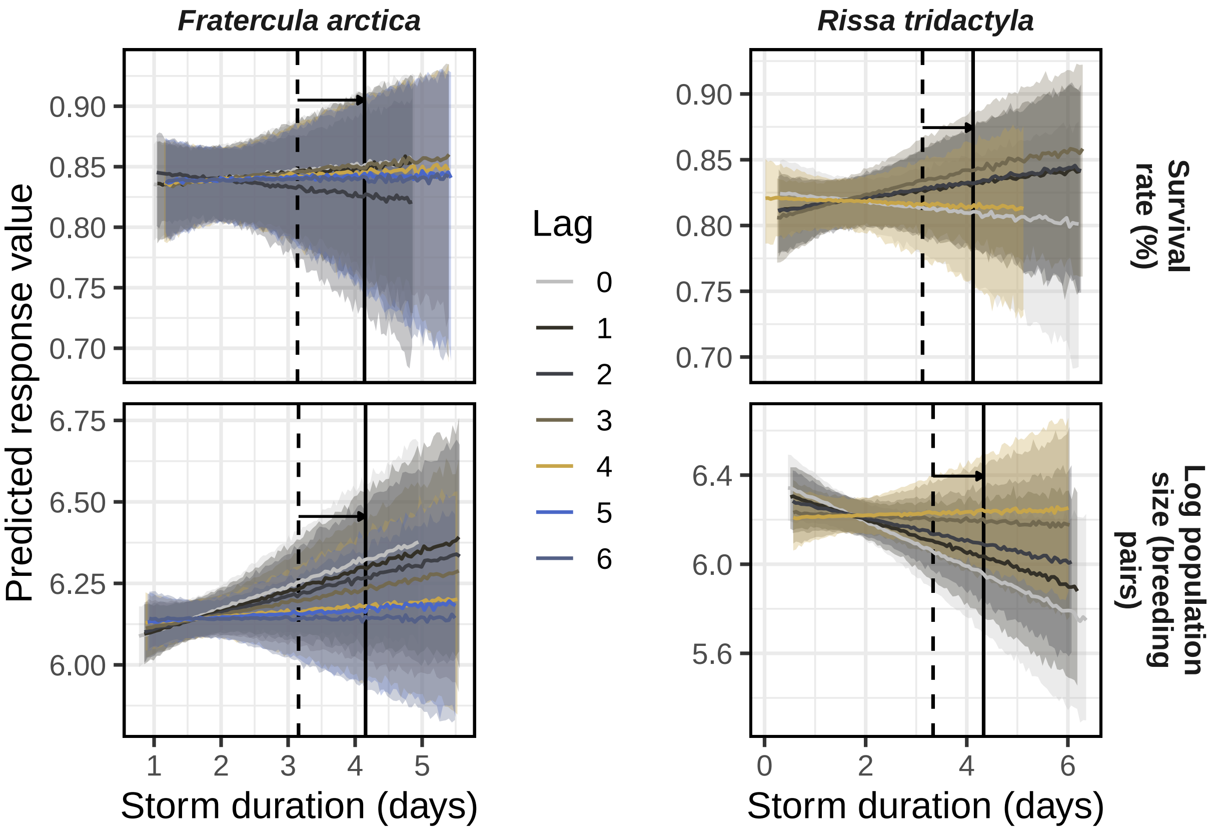


Figure S13: Marginal curves across the range of storm durations observed. Arrows indicate the predictions that inform the average treatment effect.

### S4: Supplementary tables

Table S1: Seabird population measurements and their units.

| Species | Colony | Measurement | Method | Units | Sample size | Coverage |
|---|---|---|---|---|---|---|
| *Fratercula arctica* | Anda | Population counts | Apparently occupied burrows | Number of pairs | 18 yrs | 2005-2023 |
| *Fratercula arctica* | Anda | Breeding success | Sample nest checks | Number of chicks at 20 days divided by nests | 16 yrs | 2006-2023 |
| *Fratercula arctica* | Anda | Morphometrics | Individual measurements during capture | Mass (g) & skull length (mm) | 1849 records | 2005-2024 |
| *Fratercula arctica* | Anda | Survival | Ring resightings | Binary observed-unobserved | 595 inds | 2005-2023 |
| *Fratercula arctica* | Anda | Behaviour | Global location sensors | Light intensity (lux) & conductivity (proportion of time underwater) | 21101 records | 2014-2023 |
| *Fratercula arctica* | Hjelmsøya | Population counts | Apparently occupied burrows | Number of pairs | 15 yrs | 2010-2024 |
| *Fratercula arctica* | Hjelmsøya | Breeding success | Sample nest checks | Number large chicks divided by eggs laid | 15 yrs | 2006-2023 |
| *Fratercula arctica* | Hjelmsøya | Morphometrics | Individual measurements during capture | Mass (g) & skull length (mm) | 813 records | 2004-2024 |
| *Fratercula arctica* | Hjelmsøya | Survival | Ring resightings | Binary observed-unobserved | 621 inds | 2004-2023 |
| *Fratercula arctica* | Hjelmsøya | Behaviour | Global location sensors | Light intensity (lux) & conductivity (proportion of time underwater) | 22582 records | 2014-2023 |
| *Fratercula arctica* | Hornøya | Population counts | Apparently occupied burrows | Number of pairs | 42 yrs | 1980-2024 |
| *Fratercula arctica* | Hornøya | Breeding success | Sample nest checks | Number medium chicks divided by eggs laid | 32 yrs | 1988-2023 |
| *Fratercula arctica* | Hornøya | Morphometrics | Individual measurements during capture | Mass (g) & skull length (mm) | 396 records | 2004-2021 |
| *Fratercula arctica* | Hornøya | Survival | Ring resightings | Binary observed-unobserved | 399 inds | 2004-2022 |
| *Fratercula arctica* | Hornøya | Behaviour | Global location sensors | Light intensity (lux) & conductivity (proportion of time underwater) | 50571 records | 2012-2021 |
| *Fratercula arctica* | Røst | Population counts | Apparently occupied burrows | Number of pairs | 42 yrs | 1983-2024 |
| *Fratercula arctica* | Røst | Breeding success | Sample nest checks | Number chicks fledged per egg hatched | 46 yrs | 1978-2023 |
| *Fratercula arctica* | Røst | Morphometrics | Individual measurements during capture | Mass (g) & skull length (mm) | 440 records | 1990-2023 |

Table S1: Seabird population measurements and their units.

| Species | Colony | Measurement | Method | Units | Sample size | Coverage |
|---|---|---|---|---|---|---|
| *Fratercula arctica* | Røst | Survival | Ring resightings | Binary observed-unobserved | 577 inds | 1990-2023 |
| *Fratercula arctica* | Røst | Behaviour | Global location sensors | Light intensity (lux) & conductivity (proportion of time underwater) | 56953 records | 2013-2023 |
| *Fratercula arctica* | Runde | Population counts | Apparently occupied burrows | Number of pairs | 37 yrs | 1980-2023 |
| *Fratercula arctica* | Runde | Breeding success | Sample nest checks | Number chicks fledged per egg hatched | 13 yrs | 2011-2023 |
| *Fratercula arctica* | Runde | Morphometrics | Individual measurements during capture | Mass (g) & skull length (mm) | 257 records | 2007-2024 |
| *Fratercula arctica* | Runde | Survival | Ring resightings | Binary observed-unobserved | 535 inds | 2007-2023 |
| *Fratercula arctica* | Runde | Behaviour | Global location sensors | Light intensity (lux) & conductivity (proportion of time underwater) | 22825 records | 2014-2023 |
| *Rissa tridactyla* | Ålesund | Population counts | Apparently occupied nests | Number of pairs | 13 yrs | 2011-2023 |
| *Rissa tridactyla* | Ålesund | Breeding success | Sample nest checks | Number large chicks divided by nests | 13 yrs | 2011-2023 |
| *Rissa tridactyla* | Ålesund | Morphometrics | Individual measurements during capture | Mass (g) & skull length (mm) | 210 records | 2015-2022 |
| *Rissa tridactyla* | Ålesund | Survival | Ring resightings | Binary observed-unobserved | 374 inds | 2011-2023 |
| *Rissa tridactyla* | Ålesund | Behaviour | Global location sensors | Light intensity (lux) & conductivity (proportion of time underwater) | 39443 records | 2015-2023 |
| *Rissa tridactyla* | Anda | Population counts | Apparently occupied nests | Number of pairs | 20 yrs | 2005-2024 |
| *Rissa tridactyla* | Anda | Breeding success | Sample nest checks | Number large chicks divided by nests | 17 yrs | 2006-2023 |
| *Rissa tridactyla* | Anda | Morphometrics | Individual measurements during capture | Mass (g) & skull length (mm) | 1469 records | 2005-2024 |
| *Rissa tridactyla* | Anda | Survival | Ring resightings | Binary observed-unobserved | 607 inds | 2005-2023 |
| *Rissa tridactyla* | Anda | Behaviour | Global location sensors | Light intensity (lux) & conductivity (proportion of time underwater) | 71960 records | 2009-2023 |
| *Rissa tridactyla* | Bjørnøya | Population counts | Apparently occupied nests | Number of pairs | 35 yrs | 1988-2024 |
| *Rissa tridactyla* | Bjørnøya | Breeding success | Sample nest checks | Number large chicks divided by nests | 16 yrs | 2005-2023 |
| *Rissa tridactyla* | Bjørnøya | Morphometrics | Individual measurements during capture | Mass (g) & skull length (mm) | 487 records | 2004-2024 |

Table S1: Seabird population measurements and their units.

| Species | Colony | Measurement | Method | Units | Sample size | Coverage |
|---|---|---|---|---|---|---|
| *Rissa tridactyla* | Bjørnøya | Survival | Ring resightings | Binary observed-unobserved | 429 inds | 2005-2023 |
| *Rissa tridactyla* | Bjørnøya | Behaviour | Global location sensors | Light intensity (lux) & conductivity (proportion of time underwater) | 59517 records | 2009-2023 |
| *Rissa tridactyla* | Hjelmsøya | Population counts | Apparently occupied nests | Number of pairs | 33 yrs | 1992-2024 |
| *Rissa tridactyla* | Hjelmsøya | Breeding success | Sample nest checks | Number large chicks divided by eggs laid | 15 yrs | 2006-2023 |
| *Rissa tridactyla* | Hjelmsøya | Morphometrics | Individual measurements during capture | Mass (g) & skull length (mm) | 519 records | 2004-2019 |
| *Rissa tridactyla* | Hjelmsøya | Survival | Ring resightings | Binary observed-unobserved | 470 inds | 2004-2023 |
| *Rissa tridactyla* | Hjelmsøya | Behaviour | Global location sensors | Light intensity (lux) & conductivity (proportion of time underwater) | 2868 records | 2009-2015 |
| *Rissa tridactyla* | Hornøya | Population counts | Apparently occupied nests | Number of pairs | 43 yrs | 1980-2024 |
| *Rissa tridactyla* | Hornøya | Breeding success | Sample nest checks | Number large chicks divided by nests | 36 yrs | 1988-2023 |
| *Rissa tridactyla* | Hornøya | Morphometrics | Individual measurements during capture | Mass (g) & skull length (mm) | 517 records | 2004-2020 |
| *Rissa tridactyla* | Hornøya | Survival | Ring resightings | Binary observed-unobserved | 555 inds | 2004-2021 |
| *Rissa tridactyla* | Hornøya | Behaviour | Global location sensors | Light intensity (lux) & conductivity (proportion of time underwater) | 48998 records | 2012-2023 |
| *Rissa tridactyla* | Isfjorden | Population counts | Apparently occupied nests | Number of pairs | 23 yrs | 1988-2024 |
| *Rissa tridactyla* | Isfjorden | Breeding success | Sample nest checks | Number chicks > 15 days divided by nests | 14 yrs | 2008-2023 |
| *Rissa tridactyla* | Isfjorden | Morphometrics | Individual measurements during capture | Mass (g) & skull length (mm) | 1114 records | 2008-2024 |
| *Rissa tridactyla* | Isfjorden | Survival | Ring resightings | Binary observed-unobserved | 265 inds | 2005-2023 |
| *Rissa tridactyla* | Isfjorden | Behaviour | Global location sensors | Light intensity (lux) & conductivity (proportion of time underwater) | 76633 records | 2009-2023 |
| *Rissa tridactyla* | Røst | Population counts | Apparently occupied nests | Number of pairs | 37 yrs | 1988-2024 |
| *Rissa tridactyla* | Røst | Breeding success | Sample nest checks | Number large chicks divided by nests | 37 yrs | 1988-2024 |
| *Rissa tridactyla* | Røst | Morphometrics | Individual measurements during capture | Mass (g) & skull length (mm) | 1056 records | 2005-2024 |

Table S1: Seabird population measurements and their units.

| Species | Colony | Measurement | Method | Units | Sample size | Coverage |
|---|---|---|---|---|---|---|
| *Rissa tridactyla* | Røst | Survival | Ring resightings | Binary observed-unobserved | 544 inds | 2003-2023 |
| *Rissa tridactyla* | Røst | Behaviour | Global location sensors | Light intensity (lux) & conductivity (proportion of time underwater) | 85978 records | 2008-2023 |
| *Rissa tridactyla* | Sør-Gjæslingan | Population counts | Apparently occupied nests | Number of pairs | 14 yrs | 2011-2024 |
| *Rissa tridactyla* | Sør-Gjæslingan | Breeding success | Sample nest checks | Number large chicks divided by nests | 10 yrs | 2012-2023 |
| *Rissa tridactyla* | Sør-Gjæslingan | Morphometrics | Individual measurements during capture | Mass (g) & skull length (mm) | 631 records | 2011-2024 |
| *Rissa tridactyla* | Sør-Gjæslingan | Survival | Ring resightings | Binary observed-unobserved | 335 inds | 2011-2023 |
| *Rissa tridactyla* | Sør-Gjæslingan | Behaviour | Global location sensors | Light intensity (lux) & conductivity (proportion of time underwater) | 58334 records | 2013-2023 |

Table S2: Causal estimates for a 1 magnitude increase in mean sea surface temperature. CI indicate 95% credible intervals, while Pr(x>ROPE) is the probability the estimate is larger than 1% (outside the Region Of Practical Equivalence).

| Species | Lag | Response | Driver | Median | Lower CI | Upper CI | Pr(x>ROPE) | Weight |
|---|---|---|---|---|---|---|---|---|
| *Fratercula arctica* | 0 | Foraging effort (%) | SST | -2.21 | -6.51 | 2.83 | 0.10 | 0.18 |
| *Fratercula arctica* | 1 | Foraging effort (%) | SST | 1.82 | -2.89 | 6.98 | 0.62 | 0.00 |
| *Fratercula arctica* | 2 | Foraging effort (%) | SST | -1.76 | -5.89 | 2.79 | 0.11 | 0.82 |
| *Fratercula arctica* | 3 | Foraging effort (%) | SST | -2.38 | -6.11 | 1.76 | 0.05 | 0.00 |
| *Fratercula arctica* | 4 | Foraging effort (%) | SST | -1.22 | -4.91 | 2.96 | 0.14 | 0.00 |
| *Fratercula arctica* | 5 | Foraging effort (%) | SST | -1.23 | -4.86 | 2.73 | 0.13 | 0.00 |
| *Fratercula arctica* | 6 | Foraging effort (%) | SST | 0.76 | -2.88 | 4.96 | 0.45 | 0.00 |
| *Fratercula arctica* | 0 | Mass (g) | SST | -0.40 | -4.78 | 4.22 | 0.40 | 0.00 |
| *Fratercula arctica* | 1 | Mass (g) | SST | 0.77 | -3.93 | 5.34 | 0.23 | 0.00 |
| *Fratercula arctica* | 2 | Mass (g) | SST | 0.64 | -4.26 | 5.25 | 0.25 | 0.00 |
| *Fratercula arctica* | 3 | Mass (g) | SST | 1.18 | -3.73 | 5.99 | 0.19 | 0.00 |
| *Fratercula arctica* | 4 | Mass (g) | SST | 0.84 | -3.79 | 5.54 | 0.22 | 0.00 |
| *Fratercula arctica* | 5 | Mass (g) | SST | -2.01 | -6.51 | 2.65 | 0.66 | 1.00 |
| *Fratercula arctica* | 6 | Mass (g) | SST | -0.44 | -5.26 | 4.27 | 0.40 | 0.00 |
| *Fratercula arctica* | 0 | Breeding success (%) | SST | -6.33 | -19.21 | 10.15 | 0.72 | 0.00 |
| *Fratercula arctica* | 1 | Breeding success (%) | SST | -7.80 | -18.84 | 7.65 | 0.78 | 0.00 |
| *Fratercula arctica* | 2 | Breeding success (%) | SST | -5.22 | -18.20 | 9.90 | 0.68 | 0.00 |
| *Fratercula arctica* | 3 | Breeding success (%) | SST | -17.09 | -22.55 | -3.05 | 0.99 | 0.00 |
| *Fratercula arctica* | 4 | Breeding success (%) | SST | -19.31 | -23.39 | -9.07 | 1.00 | 0.09 |
| *Fratercula arctica* | 5 | Breeding success (%) | SST | -21.49 | -24.81 | -16.25 | 1.00 | 0.45 |
| *Fratercula arctica* | 6 | Breeding success (%) | SST | -20.98 | -24.34 | -14.70 | 1.00 | 0.46 |
| *Fratercula arctica* | 0 | Survival rate (%) | SST | -0.10 | -6.22 | 4.85 | 0.38 | 0.00 |
| *Fratercula arctica* | 1 | Survival rate (%) | SST | 0.59 | -5.81 | 5.58 | 0.29 | 0.00 |
| *Fratercula arctica* | 2 | Survival rate (%) | SST | 1.02 | -4.84 | 5.82 | 0.23 | 0.00 |
| *Fratercula arctica* | 3 | Survival rate (%) | SST | 1.71 | -3.89 | 6.49 | 0.14 | 0.18 |
| *Fratercula arctica* | 4 | Survival rate (%) | SST | 0.35 | -5.38 | 5.17 | 0.30 | 0.00 |
| *Fratercula arctica* | 5 | Survival rate (%) | SST | -0.78 | -6.82 | 4.08 | 0.47 | 0.00 |
| *Fratercula arctica* | 6 | Survival rate (%) | SST | -3.97 | -9.45 | 1.69 | 0.83 | 0.82 |

Table S2: Causal estimates for a 1 magnitude increase in mean sea surface temperature. CI indicate 95% credible intervals, while Pr(x>ROPE) is the probability the estimate is larger than 1% (outside the Region Of Practical Equivalence).

| Species | Lag | Response | Driver | Median | Lower CI | Upper CI | Pr(x>ROPE) | Weight |
|---|---|---|---|---|---|---|---|---|
| *Fratercula arctica* | 0 | Population size (%) | SST | -26.12 | -41.52 | -5.86 | 0.99 | 0.00 |
| *Fratercula arctica* | 1 | Population size (%) | SST | -27.48 | -42.60 | -7.62 | 1.00 | 0.25 |
| *Fratercula arctica* | 2 | Population size (%) | SST | -31.59 | -45.82 | -13.98 | 1.00 | 0.00 |
| *Fratercula arctica* | 3 | Population size (%) | SST | -24.03 | -39.61 | -4.65 | 0.99 | 0.07 |
| *Fratercula arctica* | 4 | Population size (%) | SST | -31.85 | -45.78 | -15.20 | 1.00 | 0.03 |
| *Fratercula arctica* | 5 | Population size (%) | SST | -31.70 | -44.88 | -15.18 | 1.00 | 0.46 |
| *Fratercula arctica* | 6 | Population size (%) | SST | -27.75 | -41.56 | -9.47 | 1.00 | 0.20 |
| *Rissa tridactyla* | 0 | Foraging effort (%) | SST | -0.93 | -3.86 | 2.00 | 0.10 | 0.00 |
| *Rissa tridactyla* | 1 | Foraging effort (%) | SST | -2.07 | -4.73 | 0.88 | 0.02 | 0.89 |
| *Rissa tridactyla* | 2 | Foraging effort (%) | SST | -1.87 | -4.48 | 0.69 | 0.01 | 0.00 |
| *Rissa tridactyla* | 3 | Foraging effort (%) | SST | -2.44 | -4.87 | 0.16 | 0.01 | 0.11 |
| *Rissa tridactyla* | 4 | Foraging effort (%) | SST | -1.66 | -4.11 | 1.28 | 0.04 | 0.00 |
| *Rissa tridactyla* | 0 | Mass (g) | SST | -0.17 | -2.55 | 2.14 | 0.25 | 0.01 |
| *Rissa tridactyla* | 1 | Mass (g) | SST | 0.92 | -1.52 | 3.32 | 0.06 | 0.99 |
| *Rissa tridactyla* | 2 | Mass (g) | SST | 0.29 | -1.98 | 2.49 | 0.14 | 0.00 |
| *Rissa tridactyla* | 3 | Mass (g) | SST | -0.33 | -2.60 | 1.96 | 0.27 | 0.00 |
| *Rissa tridactyla* | 4 | Mass (g) | SST | -0.04 | -2.20 | 2.07 | 0.19 | 0.00 |
| *Rissa tridactyla* | 0 | Breeding success (%) | SST | -8.52 | -12.37 | -2.37 | 0.99 | 0.39 |
| *Rissa tridactyla* | 1 | Breeding success (%) | SST | -4.31 | -9.34 | 3.07 | 0.83 | 0.33 |
| *Rissa tridactyla* | 2 | Breeding success (%) | SST | -4.78 | -10.25 | 2.74 | 0.85 | 0.00 |
| *Rissa tridactyla* | 3 | Breeding success (%) | SST | -7.55 | -11.79 | -1.13 | 0.98 | 0.00 |
| *Rissa tridactyla* | 4 | Breeding success (%) | SST | -7.22 | -11.39 | -0.66 | 0.97 | 0.29 |
| *Rissa tridactyla* | 0 | Survival rate (%) | SST | -1.79 | -6.71 | 2.85 | 0.63 | 1.00 |
| *Rissa tridactyla* | 1 | Survival rate (%) | SST | -0.26 | -5.23 | 4.35 | 0.38 | 0.00 |
| *Rissa tridactyla* | 2 | Survival rate (%) | SST | 0.40 | -4.63 | 5.03 | 0.28 | 0.00 |
| *Rissa tridactyla* | 3 | Survival rate (%) | SST | 0.76 | -4.20 | 5.10 | 0.22 | 0.00 |
| *Rissa tridactyla* | 4 | Survival rate (%) | SST | -0.53 | -5.27 | 3.67 | 0.42 | 0.00 |
| *Rissa tridactyla* | 0 | Population size (%) | SST | -28.18 | -43.19 | -8.94 | 1.00 | 0.28 |

Table S2: Causal estimates for a 1 magnitude increase in mean sea surface temperature. CI indicate 95% credible intervals, while Pr(x>ROPE) is the probability the estimate is larger than 1% (outside the Region Of Practical Equivalence).

| Species | Lag | Response | Driver | Median | Lower CI | Upper CI | Pr(x>ROPE) | Weight |
|---|---|---|---|---|---|---|---|---|
| *Rissa tridactyla* | 1 | Population size (%) | SST | -28.37 | -42.97 | -9.96 | 1.00 | 0.42 |
| *Rissa tridactyla* | 2 | Population size (%) | SST | -32.08 | -46.23 | -14.52 | 1.00 | 0.10 |
| *Rissa tridactyla* | 3 | Population size (%) | SST | -32.25 | -45.75 | -15.60 | 1.00 | 0.18 |
| *Rissa tridactyla* | 4 | Population size (%) | SST | -32.57 | -46.43 | -15.59 | 1.00 | 0.02 |

Table S3: Causal estimates for a 1 magnitude increase in fishing pressure. CI indicate 95% credible intervals, while Pr(x>ROPE) is the probability the estimate is larger than 1% (outside the Region Of Practical Equivalence)

| Species | Lag | Response | Driver | Median | Lower CI | Upper CI | Pr(x>SESOI) | Weight |
|---|---|---|---|---|---|---|---|---|
| *Fratercula arctica* | 0 | Foraging effort (%) | Fishing pressure | 1.16 | -0.30 | 2.71 | 0.59 | 0.26 |
| *Fratercula arctica* | 1 | Foraging effort (%) | Fishing pressure | 1.10 | -0.21 | 2.59 | 0.56 | 0.41 |
| *Fratercula arctica* | 2 | Foraging effort (%) | Fishing pressure | 0.89 | -0.41 | 2.25 | 0.44 | 0.00 |
| *Fratercula arctica* | 3 | Foraging effort (%) | Fishing pressure | -0.03 | -1.30 | 1.43 | 0.07 | 0.00 |
| *Fratercula arctica* | 4 | Foraging effort (%) | Fishing pressure | -0.46 | -1.76 | 0.92 | 0.02 | 0.00 |
| *Fratercula arctica* | 5 | Foraging effort (%) | Fishing pressure | -0.40 | -1.71 | 1.04 | 0.03 | 0.00 |
| *Fratercula arctica* | 6 | Foraging effort (%) | Fishing pressure | -1.03 | -2.24 | 0.30 | 0.00 | 0.33 |
| *Fratercula arctica* | 0 | Mass (g) | Fishing pressure | -0.13 | -1.28 | 0.94 | 0.06 | 0.00 |
| *Fratercula arctica* | 1 | Mass (g) | Fishing pressure | -0.31 | -1.30 | 0.66 | 0.09 | 0.00 |
| *Fratercula arctica* | 2 | Mass (g) | Fishing pressure | -0.54 | -1.44 | 0.36 | 0.16 | 1.00 |
| *Fratercula arctica* | 3 | Mass (g) | Fishing pressure | -0.04 | -0.93 | 0.82 | 0.02 | 0.00 |
| *Fratercula arctica* | 4 | Mass (g) | Fishing pressure | 0.09 | -0.75 | 0.92 | 0.01 | 0.00 |
| *Fratercula arctica* | 5 | Mass (g) | Fishing pressure | 0.09 | -0.74 | 0.95 | 0.01 | 0.00 |
| *Fratercula arctica* | 6 | Mass (g) | Fishing pressure | 0.22 | -0.57 | 1.03 | 0.00 | 0.00 |
| *Fratercula arctica* | 0 | Breeding success (%) | Fishing pressure | -1.65 | -4.89 | 1.53 | 0.65 | 0.54 |
| *Fratercula arctica* | 1 | Breeding success (%) | Fishing pressure | -1.31 | -4.43 | 1.79 | 0.58 | 0.00 |
| *Fratercula arctica* | 2 | Breeding success (%) | Fishing pressure | -1.10 | -4.29 | 2.09 | 0.52 | 0.00 |
| *Fratercula arctica* | 3 | Breeding success (%) | Fishing pressure | 0.31 | -2.81 | 3.43 | 0.21 | 0.00 |
| *Fratercula arctica* | 4 | Breeding success (%) | Fishing pressure | 0.43 | -2.48 | 3.56 | 0.17 | 0.00 |
| *Fratercula arctica* | 5 | Breeding success (%) | Fishing pressure | -0.16 | -2.93 | 2.78 | 0.28 | 0.00 |
| *Fratercula arctica* | 6 | Breeding success (%) | Fishing pressure | -1.23 | -3.94 | 1.59 | 0.57 | 0.46 |
| *Fratercula arctica* | 0 | Survival rate (%) | Fishing pressure | -0.35 | -1.78 | 1.12 | 0.19 | 0.00 |
| *Fratercula arctica* | 1 | Survival rate (%) | Fishing pressure | 0.05 | -1.45 | 1.43 | 0.08 | 0.00 |
| *Fratercula arctica* | 2 | Survival rate (%) | Fishing pressure | 0.47 | -0.91 | 1.75 | 0.02 | 0.08 |
| *Fratercula arctica* | 3 | Survival rate (%) | Fishing pressure | -0.04 | -1.50 | 1.16 | 0.09 | 0.00 |
| *Fratercula arctica* | 4 | Survival rate (%) | Fishing pressure | 0.29 | -0.93 | 1.39 | 0.02 | 0.00 |
| *Fratercula arctica* | 5 | Survival rate (%) | Fishing pressure | 0.64 | -0.58 | 1.68 | 0.01 | 0.92 |
| *Fratercula arctica* | 6 | Survival rate (%) | Fishing pressure | 0.28 | -0.95 | 1.40 | 0.02 | 0.00 |

Table S3: Causal estimates for a 1 magnitude increase in fishing pressure. CI indicate 95% credible intervals, while Pr(x>ROPE) is the probability the estimate is larger than 1% (outside the Region Of Practical Equivalence)

| Species | Lag | Response | Driver | Median | Lower CI | Upper CI | Pr(x>SESOI) | Weight |
|---|---|---|---|---|---|---|---|---|
| *Fratercula arctica* | 0 | Population size (%) | Fishing pressure | -0.85 | -4.56 | 3.26 | 0.46 | 0.00 |
| *Fratercula arctica* | 1 | Population size (%) | Fishing pressure | -1.69 | -5.18 | 2.14 | 0.64 | 0.00 |
| *Fratercula arctica* | 2 | Population size (%) | Fishing pressure | -3.54 | -6.92 | -0.01 | 0.92 | 0.00 |
| *Fratercula arctica* | 3 | Population size (%) | Fishing pressure | -3.92 | -7.13 | -0.61 | 0.95 | 0.00 |
| *Fratercula arctica* | 4 | Population size (%) | Fishing pressure | -5.32 | -8.23 | -2.30 | 1.00 | 0.00 |
| *Fratercula arctica* | 5 | Population size (%) | Fishing pressure | -5.58 | -8.45 | -2.69 | 1.00 | 0.00 |
| *Fratercula arctica* | 6 | Population size (%) | Fishing pressure | -6.81 | -9.38 | -4.14 | 1.00 | 1.00 |
| *Rissa tridactyla* | 0 | Foraging effort (%) | Fishing pressure | 0.61 | -0.71 | 2.03 | 0.29 | 0.00 |
| *Rissa tridactyla* | 1 | Foraging effort (%) | Fishing pressure | 0.82 | -0.48 | 2.14 | 0.40 | 1.00 |
| *Rissa tridactyla* | 2 | Foraging effort (%) | Fishing pressure | 0.52 | -0.72 | 1.79 | 0.22 | 0.00 |
| *Rissa tridactyla* | 3 | Foraging effort (%) | Fishing pressure | 0.16 | -0.87 | 1.20 | 0.06 | 0.00 |
| *Rissa tridactyla* | 4 | Foraging effort (%) | Fishing pressure | 0.19 | -0.81 | 1.24 | 0.06 | 0.00 |
| *Rissa tridactyla* | 0 | Mass (g) | Fishing pressure | -0.12 | -0.88 | 0.60 | 0.01 | 0.00 |
| *Rissa tridactyla* | 1 | Mass (g) | Fishing pressure | 0.09 | -0.64 | 0.82 | 0.00 | 0.00 |
| *Rissa tridactyla* | 2 | Mass (g) | Fishing pressure | 0.04 | -0.73 | 0.78 | 0.00 | 0.00 |
| *Rissa tridactyla* | 3 | Mass (g) | Fishing pressure | 0.00 | -0.72 | 0.70 | 0.00 | 0.00 |
| *Rissa tridactyla* | 4 | Mass (g) | Fishing pressure | 0.18 | -0.57 | 0.92 | 0.00 | 1.00 |
| *Rissa tridactyla* | 0 | Breeding success (%) | Fishing pressure | -1.36 | -3.04 | 0.39 | 0.67 | 0.00 |
| *Rissa tridactyla* | 1 | Breeding success (%) | Fishing pressure | -2.40 | -3.89 | -0.90 | 0.97 | 0.52 |
| *Rissa tridactyla* | 2 | Breeding success (%) | Fishing pressure | -2.64 | -4.25 | -1.01 | 0.98 | 0.00 |
| *Rissa tridactyla* | 3 | Breeding success (%) | Fishing pressure | -2.14 | -3.83 | -0.54 | 0.92 | 0.00 |
| *Rissa tridactyla* | 4 | Breeding success (%) | Fishing pressure | -2.60 | -4.10 | -1.06 | 0.98 | 0.48 |
| *Rissa tridactyla* | 0 | Survival rate (%) | Fishing pressure | -0.52 | -1.70 | 0.55 | 0.20 | 0.01 |
| *Rissa tridactyla* | 1 | Survival rate (%) | Fishing pressure | -0.65 | -1.81 | 0.38 | 0.26 | 0.86 |
| *Rissa tridactyla* | 2 | Survival rate (%) | Fishing pressure | -0.09 | -1.19 | 0.93 | 0.05 | 0.00 |
| *Rissa tridactyla* | 3 | Survival rate (%) | Fishing pressure | -0.50 | -1.57 | 0.54 | 0.18 | 0.13 |
| *Rissa tridactyla* | 4 | Survival rate (%) | Fishing pressure | -0.30 | -1.37 | 0.71 | 0.10 | 0.00 |
| *Rissa tridactyla* | 0 | Population size (%) | Fishing pressure | -4.96 | -9.99 | 0.30 | 0.92 | 0.00 |

Table S3: Causal estimates for a 1 magnitude increase in fishing pressure. CI indicate 95% credible intervals, while Pr(x>ROPE) is the probability the estimate is larger than 1% (outside the Region Of Practical Equivalence)

| Species | Lag | Response | Driver | Median | Lower CI | Upper CI | Pr(x>SESOI) | Weight |
|---|---|---|---|---|---|---|---|---|
| *Rissa tridactyla* | 1 | Population size (%) | Fishing pressure | -6.97 | -11.48 | -1.90 | 0.99 | 0.00 |
| *Rissa tridactyla* | 2 | Population size (%) | Fishing pressure | -8.14 | -12.61 | -3.55 | 1.00 | 0.22 |
| *Rissa tridactyla* | 3 | Population size (%) | Fishing pressure | -8.18 | -12.45 | -3.74 | 1.00 | 0.06 |
| *Rissa tridactyla* | 4 | Population size (%) | Fishing pressure | -9.19 | -13.24 | -4.76 | 1.00 | 0.72 |

Table S4: Causal estimates for a 1 day increase in winter storm duration. CI indicate 95% credible intervals, while Pr(x>ROPE) is the probability the estimate is larger than 1% (outside the Region Of Practical Equivalence).

| Species | Lag | Response | Driver | Median | Lower CI | Upper CI | Pr(x>SESOI) | Weight |
|---|---|---|---|---|---|---|---|---|
| *Fratercula arctica* | 0 | Survival rate (%) | Storm duration | 0.29 | -1.33 | 1.74 | 0.06 | 0.00 |
| *Fratercula arctica* | 1 | Survival rate (%) | Storm duration | 0.22 | -1.31 | 1.67 | 0.06 | 0.00 |
| *Fratercula arctica* | 2 | Survival rate (%) | Storm duration | -0.26 | -1.80 | 1.15 | 0.17 | 0.12 |
| *Fratercula arctica* | 3 | Survival rate (%) | Storm duration | 0.23 | -1.09 | 1.54 | 0.04 | 0.54 |
| *Fratercula arctica* | 4 | Survival rate (%) | Storm duration | 0.12 | -1.34 | 1.51 | 0.06 | 0.00 |
| *Fratercula arctica* | 5 | Survival rate (%) | Storm duration | 0.07 | -1.40 | 1.47 | 0.08 | 0.00 |
| *Fratercula arctica* | 6 | Survival rate (%) | Storm duration | 0.04 | -1.45 | 1.50 | 0.08 | 0.34 |
| *Fratercula arctica* | 0 | Population size (%) | Storm duration | 3.49 | -1.04 | 8.29 | 0.03 | 0.00 |
| *Fratercula arctica* | 1 | Population size (%) | Storm duration | 2.98 | -1.16 | 7.56 | 0.03 | 0.00 |
| *Fratercula arctica* | 2 | Population size (%) | Storm duration | 2.41 | -1.87 | 6.72 | 0.06 | 0.00 |
| *Fratercula arctica* | 3 | Population size (%) | Storm duration | 1.86 | -2.11 | 6.24 | 0.08 | 0.27 |
| *Fratercula arctica* | 4 | Population size (%) | Storm duration | 0.66 | -3.34 | 4.91 | 0.20 | 0.23 |
| *Fratercula arctica* | 5 | Population size (%) | Storm duration | 0.53 | -3.35 | 4.72 | 0.22 | 0.00 |
| *Fratercula arctica* | 6 | Population size (%) | Storm duration | 0.05 | -3.90 | 4.08 | 0.30 | 0.50 |
| *Rissa tridactyla* | 0 | Survival rate (%) | Storm duration | -0.32 | -1.78 | 1.17 | 0.18 | 0.00 |
| *Rissa tridactyla* | 1 | Survival rate (%) | Storm duration | 0.45 | -1.02 | 1.94 | 0.03 | 1.00 |
| *Rissa tridactyla* | 2 | Survival rate (%) | Storm duration | 0.48 | -1.02 | 1.94 | 0.03 | 0.00 |
| *Rissa tridactyla* | 3 | Survival rate (%) | Storm duration | 0.81 | -0.82 | 2.37 | 0.01 | 0.00 |
| *Rissa tridactyla* | 4 | Survival rate (%) | Storm duration | -0.13 | -2.07 | 1.66 | 0.18 | 0.00 |
| *Rissa tridactyla* | 0 | Population size (%) | Storm duration | -7.14 | -13.35 | -0.09 | 0.96 | 0.18 |
| *Rissa tridactyla* | 1 | Population size (%) | Storm duration | -5.03 | -11.41 | 1.72 | 0.87 | 0.07 |
| *Rissa tridactyla* | 2 | Population size (%) | Storm duration | -3.44 | -9.76 | 3.17 | 0.78 | 0.09 |
| *Rissa tridactyla* | 3 | Population size (%) | Storm duration | -0.64 | -6.77 | 5.94 | 0.46 | 0.23 |
| *Rissa tridactyla* | 4 | Population size (%) | Storm duration | 0.40 | -5.88 | 6.97 | 0.34 | 0.43 |

#### Supplementary references


Alfonso, S., Gesto, M. & Sadoul, B. (2021). Temperature increase and its effects on fish stress physiology in the context of global warming. Journal of Fish Biology, 98, 1496–1508.

Bachiller, E., Skaret, G., Nøttestad, L. & Slotte, A. (2016). Feeding ecology of northeast atlantic mackerel, norwegian spring-spawning herring and blue whiting in the norwegian sea. PLOS ONE, 11, e0149238.

Barraquand, F. & Benhamou, S. (2008). Animal movements in heterogeneous landscapes: identifying profitable places and homogeneous movement bouts. Ecology, 89, 3336–3348.

Barrett, R.T. & Krasnov, Y.V. (1996). Recent responses to changes in stocks of prey species by seabirds breeding in the southern Barents Sea. ICES Journal of Marine Science, 53, 713–722.

Bråthen, V.S., Moe, B., Amélineau, F., Ekker, M., Fauchald, P., Helgason, H.H., et al. (2021). An automated procedure (v2.0) to obtain positions from light-level geolocators in large-scale tracking of seabirds. A method description for the SEATRACK project. NINA rapport. Norsk institutt for naturforskning (NINA).

Broms, C. & Melle, W. (2007). Seasonal development of Calanus finmarchicus in relation to phytoplankton bloom dynamics in the Norwegian Sea. Deep Sea Research Part II: Topical Studies in Oceanography, 54, 2760–2775.

Calenge, C. (2024). adehabitatHR: Home range estimation (manual).

Cerini, F., Childs, D.Z. & Clements, C.F. (2023). A predictive timeline of wildlife population collapse. Nature Ecology & Evolution, 7, 320–331.

Chivers, L., Lundy, M., Colhoun, K., Newton, S., Houghton, J. & Reid, N. (2012). Foraging trip time-activity budgets and reproductive success in the black-legged kittiwake. Mar Ecol Prog Ser, 456, 269–277.

Christensen-Dalsgaard, S., May, R. & Lorentsen, S.-H. (2018). Taking a trip to the shelf: Behavioral decisions are mediated by the proximity to foraging habitats in the black-legged kittiwake. Ecology and Evolution, 8, 866–878.

Collins, P.M., Green, J.A., Elliott, K.H., Shaw, P.J.A., Chivers, L., Hatch, S.A. & Halsey, L.G. (2020), Coping with the commute: behavioural responses to wind conditions in a foraging seabird. Journal of Avian Biology, 51.

Coulson, J. (2011). The Kittiwake. Poyser Monographs. 1st edn. Bloomsbury Publishing.

Cowen, R.K. & Sponaugle, S. (2009). Larval dispersal and marine population connectivity. Annual Review of Marine Science.

Cury, P.M., Boyd, I.L., Bonhommeau, S., Anker-Nilssen, T., Crawford, R.J.M., Furness, R.W., et al. (2011). Global seabird response to forage fish depletion—one-third for the birds. Science, 334, 1703–1706.

Dee, L.E., Ferraro, P.J., Severen, C.N., Kimmel, K.A., Borer, E.T., Byrnes, J.E.K., et al. (2023). Clarifying the effect of biodiversity on productivity in natural ecosystems with longitudinal data and methods for causal inference. Nature Communications, 14, 2607.

Engelhard, G. & Heino, M. (2004). Maturity changes in Norwegian spring-spawning herring Clupea harengus: Compensatory or evolutionary responses? Mar Ecol Prog Ser, 272, 245–256.

Fauchald, P., Tarroux, A., Bråthen, V.S., Descamps, S., Ekker, M., Helgason, H.H., et al. (2019). Arctic-breeding seabirds' hotspots in space and time - A methodological framework for year-round modelling of environmental niche and abundance using light-logger data ( No. NINA Report 1657). Norwegian Institute for Nature Research.

Fayet, A.L., Clucas, G.V., Anker-Nilssen, T., Syposz, M. & Hansen, E.S. (2021). Local prey shortages drive foraging costs and breeding success in a declining seabird, the Atlantic puffin. Journal of Animal Ecology, 90, 1152–1164.

Fayet, A.L., Freeman, R., Shoji, A., Boyle, D., Kirk, H.L., Dean, B.J., et al. (2016). Drivers and fitness consequences of dispersive migration in a pelagic seabird. Behavioral Ecology, 27, 1061–1072.

Fayet A.L., Freeman R., Anker-Nilssen T., Diamond A., Erikstad K.E., Fifield D., Fitzsimmons M.G., Hansen E.S., Harris M.P., Jessopp M., Kouwenberg A.L., Kress S., Mowat S., Perrins .CM., Petersen A., Petersen I.K., Reiertsen T.K., Robertson G.J., Shannon P., Sigurðsson I.A., Shoji A., Wanless S. & Guilford T. (2017) Ocean-wide drivers of migration strategies and their influence on population breeding performance in a declining seabird. Current Biology, 18, 27(24):3871-3878.e3.

Freitas, C., Villegas-Ríos, D., Moland, E. & Olsen, E.M. (2021). Sea temperature effects on depth use and habitat selection in a marine fish community. Journal of Animal Ecology, 90, 1787–1800.

Gibson, D., Riecke, T.V., Catlin, D.H., Hunt, K.L., Weithman, C.E., Koons, D.N., et al. (2023). Climate change and commercial fishing practices codetermine survival of a long-lived seabird. Global Change Biology, 29, 324–340.

Goldenberg, S.U., Taucher, J., Fernández-Méndez, M., Ludwig, A., Arístegui, J., Baumann, M., et al. (2022). Nutrient composition (Si:N) as driver of plankton communities during artificial upwelling. Frontiers in Marine Science, Volume 9-2022.

Harris, M.P. & Wanless, S. (2011). The puffin. A&C Black.

IPCC. (2023). Climate change 2021 – the physical science basis: working group I contribution to the sixth assessment report of the intergovernmental panel on climate change. Cambridge University Press, Cambridge.

Irons, D.B., Anker-Nilssen, T., Gaston, A., Byrd, G.V., Falk, K., Gilchrist, G., et al. (2008). Fluctuations in circumpolar seabird populations linked to climate oscillations. Global Change Biology, 14, 1455–1463.

Johnson, C., Inall, M. & Häkkinen, S. (2013). Declining nutrient concentrations in the northeast Atlantic as a result of a weakening Subpolar Gyre. Deep Sea Research Part I: Oceanographic Research Papers, 82, 95–107.

Jones, T., Parrish, J.K., Peterson, W.T., Bjorkstedt, E.P., Bond, N.A., Ballance, L.T., et al. (2018). Massive mortality of a planktivorous seabird in response to a marine heatwave. Geophysical Research Letters, 45, 3193–3202.

Kooijman, S.A.L.M., Sousa, T., Pecquerie, L., Van Der Meer, J. & Jager, T. (2008). From food-dependent statistics to metabolic parameters, a practical guide to the use of dynamic energy budget theory. Biological Reviews, 83, 533–552.

Kubinec, R. (2023). Ordered beta regression: a parsimonious, well-fitting model for continuous data with lower and upper bounds. Political Analysis, 31, 519–536.

Labocha, M.K. & Hayes, J.P. (2012). Morphometric indices of body condition in birds: a review. Journal of Ornithology, 153, 1–22.

Laurenson, K., Wood, M.J., Birkhead, T.R., Priestley, M.D.K., Sherley, R.B., Fayet, A.L., et al. (2025). Long-term multi-species demographic studies reveal divergent negative impacts of winter storms on seabird survival. Journal of Animal Ecology, 94, 139–153.

Lorentsen, S.H. & Christensen-Dalsgaard, S. (2009). Det nasjonale overvåkingsprogrammet for sjøfugl. Resultater til og med hekkesesongen 2008 ( No. 439). NINA.

Maunder, M.N. & Punt, A.E. (2004). Standardizing catch and effort data: a review of recent approaches. Fisheries Research, 70, 141–159.

Oro, D. & Furness, R.W. (2002). Influences of food availability and predation on survival of kittiwakes. Ecology, 83, 2516–2528.

Pelegrí, J.L., Marrero-Díaz, A. & Ratsimandresy, A.W. (2006). Nutrient irrigation of the North Atlantic. Progress in Oceanography, 70, 366–406.

Phillips, R.A., Thompson, D.R. & Hamer, K.C. (1999). The impact of great skua predation on seabird populations at St Kilda: a bioenergetics model. Journal of Applied Ecology, 36, 218–232.

Ponchon, A., Grémillet, D., Christensen-Dalsgaard, S., Erikstad, K.E., Barrett, R.T., Reiertsen, T.K., et al. (2014). When things go wrong: intra-season dynamics of breeding failure in a seabird. Ecosphere, 5, art4.

Pradel, R., Hines, J.E., Lebreton, J.-D. & Nichols, J.D. (1997). Capture-recapture survival models taking account of transients. Biometrics, 53, 60–72.

Pradel, R. & Sanz-Aguilar, A. (2012). Modeling trap-awareness and related phenomena in capture-recapture studies. PLOS ONE, 7, e32666.

Reiertsen, T.K., Erikstad, K.E., Anker-Nilssen, T., Barrett, R.T., Boulinier, T., Frederiksen, M., et al. (2014). Prey density in non-breeding areas affects adult survival of black-legged kittiwakes Rissa tridactyla. Mar Ecol Prog Ser, 509, 289–302.

Reiertsen, T.K., Layton-Matthews, K., Erikstad, K.E., Hodges, K., Ballesteros, M., Anker-Nilssen, T., et al. (2021). Inter-population synchrony in adult survival and effects of climate and extreme weather in non-breeding areas of Atlantic puffins. Mar Ecol Prog Ser, 676, 219–231.

Renner, H.M., Piatt, J.F., Renner, M., Drummond, B.A., Laufenberg, J.S. & Parrish, J.K. (2024). Catastrophic and persistent loss of common murres after a marine heatwave. Science, 386, 1272–1276.

Rose, G.A. (2005). On distributional responses of North Atlantic fish to climate change. ICES Journal of Marine Science, 62, 1360–1374.

Sandvik, H., Erikstad, K., Barrett, R. & Yoccoz, N. (2005). The effect of climate on adult survival in five species of North Atlantic seabirds. Journal of Animal Ecology, 74, 817–831.

Simmonds, E.G., Adjei, K.P., Cretois, B., Dickel, L., González-Gil, R., Laverick, J.H., et al. (2024). Recommendations for quantitative uncertainty consideration in ecology and evolution. Trends in Ecology & Evolution, 39, 328–337.

Slotte, A., Salthaug, A., Vatnehol, S., Johnsen, E., Mousing, E.A., Høines, Å., et al. (2025). Herring spawned poleward following fishery-induced collective memory loss. Nature, 642, 965–972.

Stollewerk, A., Kratina, P., Sentis, A., Chaparro-Pedraza, C., Decaestecker, E., De Meester, L., et al. (2025). Plasticity in climate change responses. Biological Reviews, n/a.

Sydeman, W.J., Schoeman, D.S., Thompson, S.A., Hoover, B.A., García-Reyes, M., Daunt, F., et al. (2021). Hemispheric asymmetry in ocean change and the productivity of ecosystem sentinels. Science, 372, 980–983.

Teplitsky, C., Mills, J.A., Alho, J.S., Yarrall, J.W. & Merilä, J. (2008). Bergmann's rule and climate change revisited: Disentangling environmental and genetic responses in a wild bird population. Proceedings of the National Academy of Sciences, 105, 13492–13496.

Thieurmel, B. & Elmarhraoui, A. (2022). suncalc: Compute sun position, sunlight phases, moon position and lunar phase (manual).

Villejo, S.J., Martino, S., Illian, J., Ryan, W. & Lindgren, F. (2025). Validating uncertainty propagation approaches for two-stage Bayesian spatial models using simulation-based calibration. arXiv e-prints, arXiv:2502.18962.

Walsh, P.M., Halley, D.J., Harris, M.P., del Nevo, A., Sim, I.M.W. & Tasker, M.L. (1995). Seabird monitoring handbook for Britain and Ireland: a compilation of methods for survey and monitoring of breeding seabirds. JNCC/RSPB/ITE/Seabird Group, Peterborough.

Wong, B.B.M. & Candolin, U. (2015). Behavioral responses to changing environments. Behavioral Ecology, 26, 665–673.

Wooldridge, J.M. (1999). Distribution-free estimation of some nonlinear panel data models. Journal of Econometrics, 90, 77–97.
